# A type 1 immune-stromal cell network mediates disease tolerance and barrier protection against intestinal infection

**DOI:** 10.1101/2024.09.04.611190

**Authors:** Susan Westfall, Maria E. Gentile, Tayla M. Olsen, Danielle Karo-Atar, Andrei Bogza, Franziska Röstel, Ryan D. Pardy, Giordano Mandato, Ghislaine Fontes, De’Broski Herbert, Heather J. Melichar, Valerie Abadie, Martin J. Richer, Donald C. Vinh, Joshua F. E. Koenig, Oliver J. Harrison, Maziar Divangahi, Sebastian Weis, Alex Gregorieff, Irah L. King

## Abstract

Type 1 immunity mediates host defense through pathogen elimination, but whether this pathway also impacts tissue function is unknown. Here we demonstrate that rapid induction of IFNγ signaling coordinates a multi-cellular response that is critical to limit tissue damage and maintain gut motility following infection of mice with a tissue-invasive helminth. IFNγ production is initiated by antigen-independent activation of lamina propria CD8^+^ T cells following MyD88-dependent recognition of the microbiota during helminth-induced barrier invasion. IFNγ acted directly on intestinal stromal cells to recruit neutrophils that limited parasite-induced tissue injury. IFNγ sensing also limited the expansion of smooth muscle actin-expressing cells to prevent pathological gut dysmotility. Importantly, this tissue-protective response had limited impact on parasite burden, indicating that IFNγ supports a disease tolerance defense strategy. Our results have important implications for managing the pathophysiological sequelae of post-infectious gut dysfunction and chronic inflammatory diseases associated with stromal remodelling.

**HIGHLIGHTS:**

1. Type 1 immunity is required for disease tolerance to tissue-invasive infection.
2. Gut-resident CD8^+^ T cells produce IFNγ in an antigen-independent, yet microbiota-dependent manner.
3. IFNγ signaling recruits neutrophils in a cell-extrinsic manner to limit helminth-induced tissue injury.
4. Direct sensing of IFNγ by intestinal stroma is essential to limit tissue damage and maintain gut motility during infection.

## INTRODUCTION

Decades of research have made incredible advances in our understanding of host resistance, that is, pathogen elimination. However, our knowledge of pathways that limit tissue damage to ensure survival in the face of persistent infection, a host defense strategy referred to as disease tolerance, is comparatively limited^1^. A relevant, yet understudied, setting to investigate the mechanisms of disease tolerance is intestinal helminth infection^2^. Helminths are large, tissue-invasive parasites that migrate through host tissues to complete their life cycle, resulting in profound tissue damage during infection. Remarkably, most enteric helminths colonize their hosts without inducing overt morbidity, likely contributing to their pervasiveness across mammalian species^3^.

Compared to microbial pathogens such as bacteria, viruses and fungi, the size and tissue-invasive nature of helminths pose unique challenges to their hosts. As a result, type 2 immune responses including production of interleukin (IL)-4 and IL-13, likely evolved to selectively promote the expulsion of diverse enteric helminth species by enhancing the physiological functions of the intestine including motility, epithelial turnover and mucus secretion^4^. Tissue-specific type 2 cytokine signaling also induces *de novo* generation of reparative monocytes/macrophages that execute wound healing^5^. However, the “weep and sweep” and tissue repair functions of type 2 immunity are dominant during the luminal stage of helminth infection, leaving unanswered fundamental questions about the host defense pathways that limit the tissue damage imposed by these parasites and retain organ function during the invasive stage of infection. Thus, understanding the mechanisms by which the host endures tissue invasion by enteric helminths provides a rich opportunity to identify pathways of resilience that may be applicable to diverse settings of barrier injury.

Investigating infection of mice using the enteric parasitic roundworm *Heligmosomoides polygyrus bakeri* (*Hpb*), we and others have described a transient, yet robust type 1 immune response during the tissue-invasive stage of infection^6–8^. This response is characterized by IFNγ-dependent transcription of interferon-stimulated genes (ISGs) that largely precedes the canonical type 2 immune response^6^. Although classically involved in microbe elimination and anti-tumor immunity^9^, ISG expression during *Hpb* infection has been suggested to promote regeneration of the gut barrier by inducing Sca-1^+^ fetal-like intestinal epithelial cells (IECs)^7^. In addition, direct IFNγ signaling on enteric glial cells acts to repair *Hpb*-induced granulomatous lesions^8^. IFNγ signaling also preserves barrier integrity by recruiting tissue-protective natural killer (NK) cells to the *Hpb* infection site^6^. These effects were independent of changes in parasite burden suggesting that, in this context, type 1 immunity may be integral to disease tolerance^1,10^.

Here we performed an in-depth analysis of the cellular sources and targets of IFNγ signaling to reveal the fundamental type I immune networks required to endure an enteric helminth infection. We demonstrate that a breach of the epithelial barrier elicits microbiota-dependent activation of tissue-resident lamina propria CD8^+^ T cells that rapidly produce IFNγ in an antigen-independent manner. Despite widespread IFNγ signaling at the site of infection, direct sensing of this cytokine by the small intestinal stromal cell compartment is critical for the recruitment of tissue-protective neutrophils and restraining expansion of smooth muscle actin-expressing cells. Indeed, germline or stromal cell-specific deletion of the IFNγ receptor resulted in enhanced granulomatous pathology and, following high dose parasite challenge, severe gut dysmotility and mortality. Collectively, this study reveals the unexpected functions of IFNγ in disease tolerance and provides a new conceptual framework for investigating how immune pathways, conventionally thought to only mediate host resistance, ensure tissue preservation during infection.

## RESULTS

### Type 1 immune activation during early *Hpb* infection controls parasite-induced tissue damage and preserves organ function

We and others previously showed that a type 1 immune response, characterized by IFNγ target gene expression, dominates the intestinal immune landscape during the tissue-invasive phase of *Hpb* infection^6,7^. To further evaluate the dynamics of IFNγ signaling, we performed a kinetic analysis of ISG expression during the tissue-invasive (days 1-6) and luminal stages (days 8-14) of *Hpb* infection. We found ISG expression to be robust yet transient with *Cxcl9, Cxcl10* and *Ly6a* (encoding for Sca-1) mRNA all highly expressed at day 2 *Hpb* infection and rapidly decreasing thereafter (Figure 1A). Similarly, surface Sca-1 expression by intestinal epithelial cells also peaked between days 2-4 (Figure 1B, C). This response was specific to IFNγ signaling because *Hpb*-infected mice with a germline deletion of the IFNγ receptor (*Ifngr^-/-^*) failed to induce ISGs at these time points compared to IFNγR-sufficient (*Ifngr*^+/+^) animals (Figure 1D). As IFNγ signaling is antagonistic to canonical type 2 immune activation^11^, the loss of ISG expression in *Hpb*-infected *Ifngr*^-/-^ mice corresponded to an enhanced type 2 immune signature compared to controls at 6-8 days post-infection (Figure S1).

**Figure 1:**
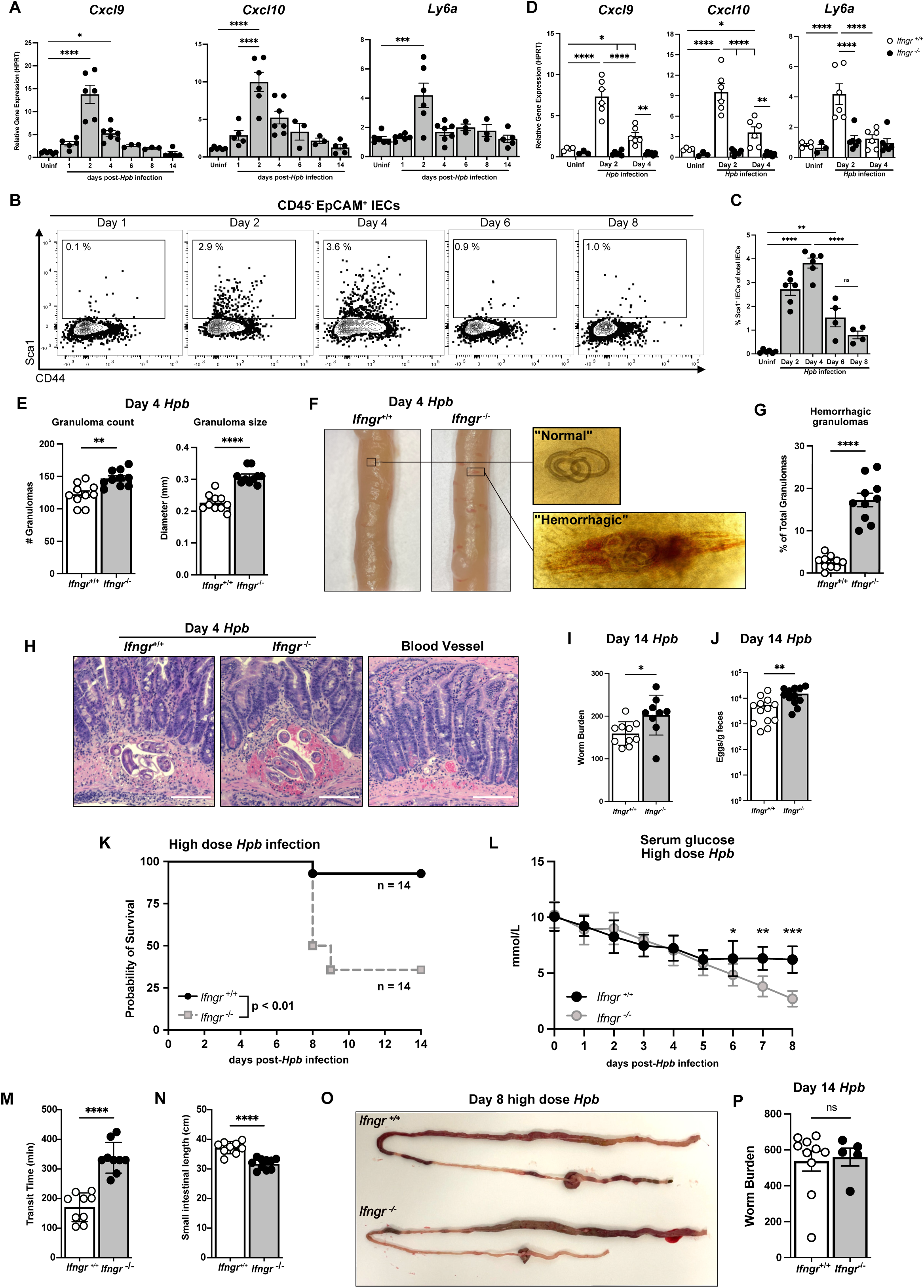
Transient IFNγ signaling limits tissue damage and mediates disease tolerance to tissue-invasive helminth infection. (A) qPCR analysis of ISG mRNA expression from whole small intestinal tissue (proximal duodenum) at the indicated time points after *Hpb* infection. (B) Representative flow cytometry contour plots of Sca-1 expression by small intestinal epithelial cells (IECs) gated on Viable^+^CD45^-^ EpCAM^+^ cells. (C) The frequency of Sca-1^+^ cells at various time points of *Hpb* infection. Each dot represents an individual mouse. Two independent pooled experiments, n=3-7 per group. (D) qPCR analysis of ISG mRNA expression from whole small intestinal tissue from *Ifngr*^+/+^ and *Ifngr^-^* ^/-^ mice at the indicated time points. Two independent pooled experiments, n=6 mice per group. (E) Enumeration of total granulomas (left) in the top 10 cm of the small intestine and their diameter (right) at day 4 *Hpb* infection. The average of 30 granulomas per mouse was used to generate each data point. (F) Representative images of the duodenum of *Ifngr*^+/+^ and *Ifngr*^-/-^ mice at day 4 *Hpb* infection with an inset of normal and “hemorrhagic” granulomas. (G) Frequency of hemorrhagic granulomas in the top 10 cm of the small intestine. Three independent experiments, n=10 mice per group. (H) Representative image of H&E staining showing bleeding in *Ifngr*^+/+^ and *Ifngr^-/-^* mice at day 4 *Hpb* infection. Scale bar, 50 µm. A blood vessel is shown for a positive control for blood staining. (I) Worm and (J) egg burden of *Hpb* infected *Ifngr*^+/+^ and *Ifngr*^-/-^ mice at day 14 *Hpb* infection. Three independent experiments, n=8-10 mice per group. (K) Kaplan Meier survival curve of high dose (600 L3 larvae) *Hpb* infected mice, three independent experiments, n=14 mice per group. (L) Systemic blood glucose levels (unfasted) in high-dose infected mice measured at the same time daily. Two independent experiments, n=16 per group. (M) Transit time and (N) small intestinal length of high dose infected mice at day 7 *Hpb* infection. Two independent pooled experiments, n=8-9 mice per group. (O) Representative image of intestines at day 7 *Hpb* infection. (P) Worm burden in surviving high dose infected mice at day 14 *Hpb* infection. Two independent pooled experiments, n=5-10 mice per group. Between group comparisons analyzed with student’s t test. For time course analyses, a 2-way ANOVA with Tukey’s posthoc analysis was performed. *p<0.05, **p<0.01, ***p<0.001, ****p<0.0001. Survival curves analyzed with log-rank (Mantel-Cox) test.

To examine the impact of IFNγ signals on host defense during the tissue-invasive stage of *Hpb* infection, we assessed gross changes to the larval-induced granulomas in IFNγR-sufficient and - deficient mice. At day 4, we observed an increase in the number and size of visible granulomas in *Ifngr*^-/-^ mice compared to controls (Fig. 1E). We also found an increased frequency of granulomatous lesions in the *Ifngr*^-/-^ mice (Figure 1F, G) which were confirmed to be hemorrhagic by a positive blood stain with intense Eosin Y signal (Figure 1H) as well as detection of intravenously administered TRITC-Dextran dye in the granulomas of *Ifngr*^-/-^, but not control, mice (Figure S2A). Conversely, treatment of *Ifngr*^+/+^ mice with low dose recombinant IFNγ decreased the number and size of visible granulomas (Figure S2B) suggesting that IFNγ reduces helminth-induced tissue remodelling. Despite the increased frequency of hemorrhagic granulomas in the absence of IFNγ signals during infection, we did not detect a decrease in barrier permeability as quantified by serum FITC-Dextran following gavage (Figure S2C) or circulating endotoxin levels (Figure S2D). Importantly, the increased tissue morbidity observed in the *Ifngr*^-/-^ mice was associated with a small, but significant increase in worm (Figure 1I) and egg burden (Figure 1J) at day 14. These results indicate that transient activation of IFNγ signals restrain helminth-induced tissue damage.

Although we observed changes in tissue damage in the absence of IFNγ signals using a conventional infection dose (200 L3 larvae), no appreciable sickness behavior (e.g. weight loss or altered posturing) was observed. To test whether a higher dose of *Hpb* larvae would result in disease manifestation, we challenged mice with 600 L3 larvae. Whereas *Ifngr*^+/+^ mice displayed minimal signs of sickness behavior at any time point examined (Figure S3A), *Ifngr*^-/-^ mice exhibited a significant reduction in systemic glucose levels and a 75% mortality rate by day 8 in high dose infected mice (Figure 1K, L). While we detectd a larger increase in serum endotoxin levels in the *Ifngr*^-/-^ mice compared to controls (Figure S3B), sepsis did not appear to be a key driver of mortality as no difference in systemic bacterial translocation was observed between groups (Figure S3C), and there was limited evidence of organ failure based on serological testing (Figure S3D) or serum cytokine quantification except for increased IFNγ, IL-17 and IL-6 in high dose infected *Ifngr*^-/-^ mice (Figure S3E). In contrast, intestinal transit time in *Ifngr*^-/-^ mice was severely compromised at 8 days after high dose *Hpb* infection compared to control animals as determined by a carmine red assay (Figure 1M), which was coupled with small intestinal distention and shortening (Figure 1N, O). Finally, the elevated mortality in *Ifngr*^-/-^ mice was not associated with a difference in parasite burden between surviving mice (Figure 1P). Collectively, these results indicate that transient IFNγ signaling promotes disease tolerance by limiting tissue damage and ensuring intestinal function during tissue invasion by an enteric helminth, independent of parasite burden control.

### Antigen-independent production of IFNγ by tissue-resident CD8^+^ T cells drives the type 1 immune signature during *Hpb* infection

Diverse subsets of immune cells are capable of producing IFNγ during early *Hpb* infection^7^. Using IFNγ reporter mice in which *Ifng* mRNA is detectable by eYFP expression^12^, we found an increase in eYFP^+^ cells as early as day 2 post-infection and confirmed that more than 90% of these cells were either TCRβ^+^ αβ T cells or NK1.1^+^ TCRβ^-^ NK cells (Figure 2A). Depletion of NK cells from the small intestinal lamina propria (SILP) during *Hpb* infection^6^ did not impact ISG induction (Figure 2B). By contrast, infection of TCRβ^-/-^ mice abrogated ISG expression compared to controls (Figure 2C). To test whether IFNγ production by T cells was sufficient to induce ISG expression, TCRβ-deficient mice received wildtype or IFNγ^-/-^ T cells and, two weeks later, were infected with *Hpb*. Notably, the transfer of wildtype, but not IFNγ-deficient, T cells restored ISG expression in TCRβ-deficient animals (Figure 2D). Characterization of the T cell subsets revealed that SILP CD8^+^ ɑβ T cells were the dominant population induced to express *Ifng* at the peak time of ISG expression (Figure 2E). Mirroring the ISG expression profile (Figure 1A), the frequency of *Ifng*-expressing eYFP^+^ CD8^+^ T cells peaked at days 2 and 4 (Figure 2F, G) and was further supported by in vitro PMA/ionomycin stimulation at the same time points in non-reporter mice (Figure S4).

**Figure 2:**
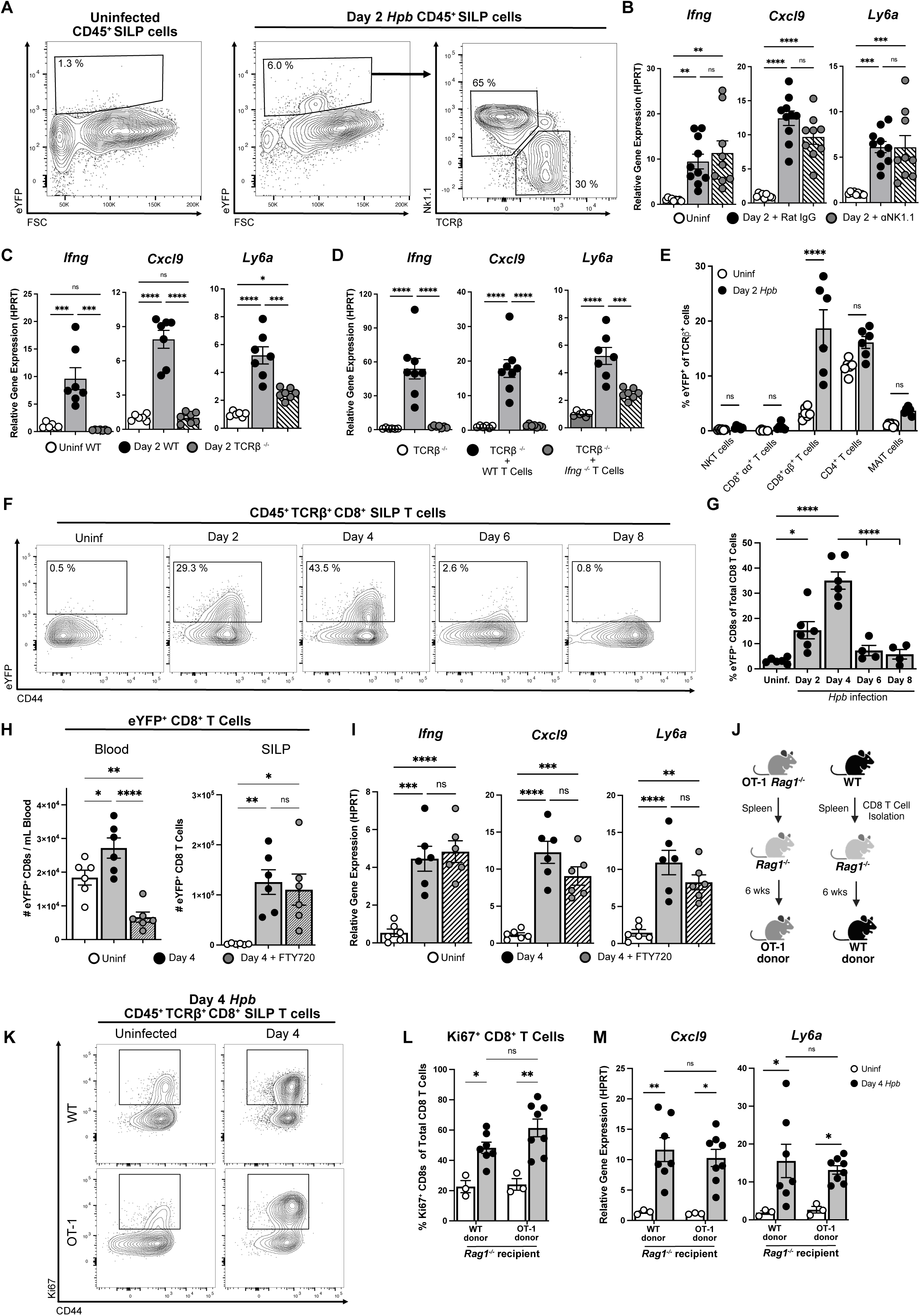
Tissue-resident CD8 T cells produce IFNγ independent of cognate antigen. (A) Representative flow cytometry plot of total eYFP^+^ cells from the SILP of uninfected and day 2 *Hpb*-infected mice, gated on viable, CD45^+^ cells. The plot on the right shows TCRβ^+^ or NK1.1^+^ IFNγ-expressing cells based on eYFP^+^ cells. (B) SILP ISG expression in WT mice treated with isotype control or anti-NK1.1 antibody at day 2 *Hpb* infection. (C) SILP ISG expression in WT or TCRβ^-/-^ mice at day 2 *Hpb*. (D) SILP ISG expression in day 2 *Hpb* TCRβ^-/-^ mice following the adoptive transfer of total T cells from either WT or *Ifng*^-/-^ mice. (A-D) All data shown represent pooled results from 2-3 independent experiments with n=8-9 mice per group. (E) IFNγ-expressing T cell subtypes detected as eYFP^+^ cells at day 2 *Hpb* infection. (F) Representative flow cytometry plots of eYFP^+^ CD44^+^ CD8 T cells during *Hpb* infection, gated on SILP viable CD45^+^ TCRβ^+^ CD8^+^ cells, quantified across multiple mice shown in (G). (H) Number of eYFP^+^ CD8 T cells in the blood (left) and SILP (right) of day 4 *Hpb* infected, FTY720-treated mice with (I) associated ISG data from the SILP. Two independent pooled experiments with n=6 mice per group. (J) Schema of OT-1 adoptive transfer experiment for results shown in K-M. (K) Representative flow cytometry plots of proliferating OT-1 derived or WT-derived CD8 T cells from day 4 *Hpb* infected mice. Gated on SILP viable CD45^+^ TCRβ^+^ CD8^+^ T cells. (L) Proliferation (Ki67^+^) of CD8^+^ T cells and (M) SILP ISG expression from two independent pooled experiments with n=3-7 mice per group. Between group comparisons analyzed with student’s t test while time course analyses with 1- or 2-way ANOVAs with Tukey’s posthoc analysis. *p<0.05, **p<0.01, ***p<0.001, and ****p<0.0001.

To determine whether the *Ifng*-expressing eYFP^+^ CD8^+^ T cells were tissue-resident or derived from the circulation, we first performed immunophenotyping. Our results indicated that eYFP^+^ cells expressed a CD44^+^CD62L^-^ memory phenotype with variable expression of CD49d, CD103, CD69, PD-1 and ICOS (Figure S5). We also intravenously administered a fluorochrome-conjugated anti-CD45 antibody into day 4 *Hpb*-infected mice 3 minutes before sacrifice, demonstrating that eYFP^+^ CD8^+^ T cells found in the SILP cell preparation were not within the intestinal vasculature (Figure S6). Notably, we could detect an increase in eYFP^+^ CD8^+^ T cells in several peripheral tissue compartments but not until day 4, a later time point than detectable CD8^+^ T cell activation in the SILP (Figure S7). Consistently, expression of the gut-homing integrin ɑ4β7 was not detectable in SILP CD8^+^ T cells until day 4, which would be expected on newly immigrated CD8^+^ T cells from circulation (Figure S8). To formally test whether CD8^+^ tissue-resident memory (Trm) T cells were critical for ISG expression in the *Hpb*-infected intestine, we blocked migration of circulating T cells by administration of the sphingosine 1-phosphate receptor agonist FTY720^13^ prior to and during infection. As expected, FTY720 treatment led to a significant decrease in the number of detectable eYFP^+^ CD8^+^ T cells in the circulation. However, FTY720 had no impact on the number of SILP eYFP^+^ CD8^+^ T cells or induction of ISG expression in the small intestine (Figure 2H, I).

Given the rapid induction of IFNγ-producing T cells during *Hpb* infection and that no *Hpb*-derived MHCI-restricted antigens have been described, we hypothesized that SILP CD8^+^ T cells were activated in an antigen-independent manner. To test this idea, we used a previously published protocol^14^ in which we transferred CD8^+^ T cells from the spleens of C57BL/6 or OT-1 x Rag1^-/-^ animals, in which the entire lymphocyte repertoire of the latter is OVA-specific, into Rag1^-/-^ recipients six weeks prior to *Hpb* infection (Figure 2J). Consistent with prior results^14^, we readily detected donor T cells from each group in the SILP of uninfected and infected recipients (Figure 2K). Furthermore, regardless of antigen specificity, we observed an equal increase in proliferating Ki67^+^ CD8^+^ T cells and ISG induction in both groups of recipients at day 4 (Figure 2K-M). In a complementary approach, we observed similar activation of Thy1.1^+^ LCMV-specific P14 donor cells in C57BL/6 mice (hosting an endogenous T cell repertoire) previously infected with LCMV-Armstrong and then challenged with *Hpb* one month later (Figure S9). Together, these results indicate that *Hpb* infection induces activation and IFNγ production by lamina propria CD8^+^ Trm T cells in an antigen-independent manner.

### IFNγ production during *Hpb* infection requires the gut microbiota and MyD88 signaling

To investigate the trigger of IFNγ production by Trm CD8^+^ T cells, we tested several antigen-independent modes of activation. The alarmins, IL-18, IL-33 and IL-36γ, although upregulated at the transcriptional level during early *Hpb* infection, were all dispensable for IFNγ production from CD8^+^ T cells (Figure S10). In addition, IL-15 neutralization, while eliminating *Hpb*-induced NK cell recruitment to the SILP, had no impact on the frequency of IFNγ^+^ CD8^+^ T cells or Sca-1 expression by IECs (Figure S11). One compartment that has been previously implicated in intestinal CD8^+^ T cell activation is the gut microbiota^15^. To directly test whether the gut microbiota was required for IFNγ production and ISG induction, we implemented our recently-developed model of germ-free (GF) *Hpb* infection^16^. To this end, GF mice were infected with 200 GF *Hpb* larvae and at day 4, IFNγ and ISG expression were assessed. Littermate GF mice reconstituted with an SPF microbiota (i.e. conventionalized mice) were used as a positive control (Figure 3A). Importantly, we have previously shown that the adult worm burden and fecundity of GF *Hpb* are comparable to conventionally-reared SPF larvae in both SPF and GF hosts^16^. Using this approach, we found that ISG and IFNγ induction was entirely dependent on the presence of the gut microbiota (Figure 3B-D). Similarly, helminth-infected GF mice phenocopied many features of infected *Ifngr*^-/-^ mice in gross tissue damage (Figure 3E), increased number, size and hemorrhaging of detectable granulomas (Figure 3F) as well as an overall shortening of the small intestine following a high dose infection, compared to conventionalized controls (Figure 3G).

**Figure 3:**
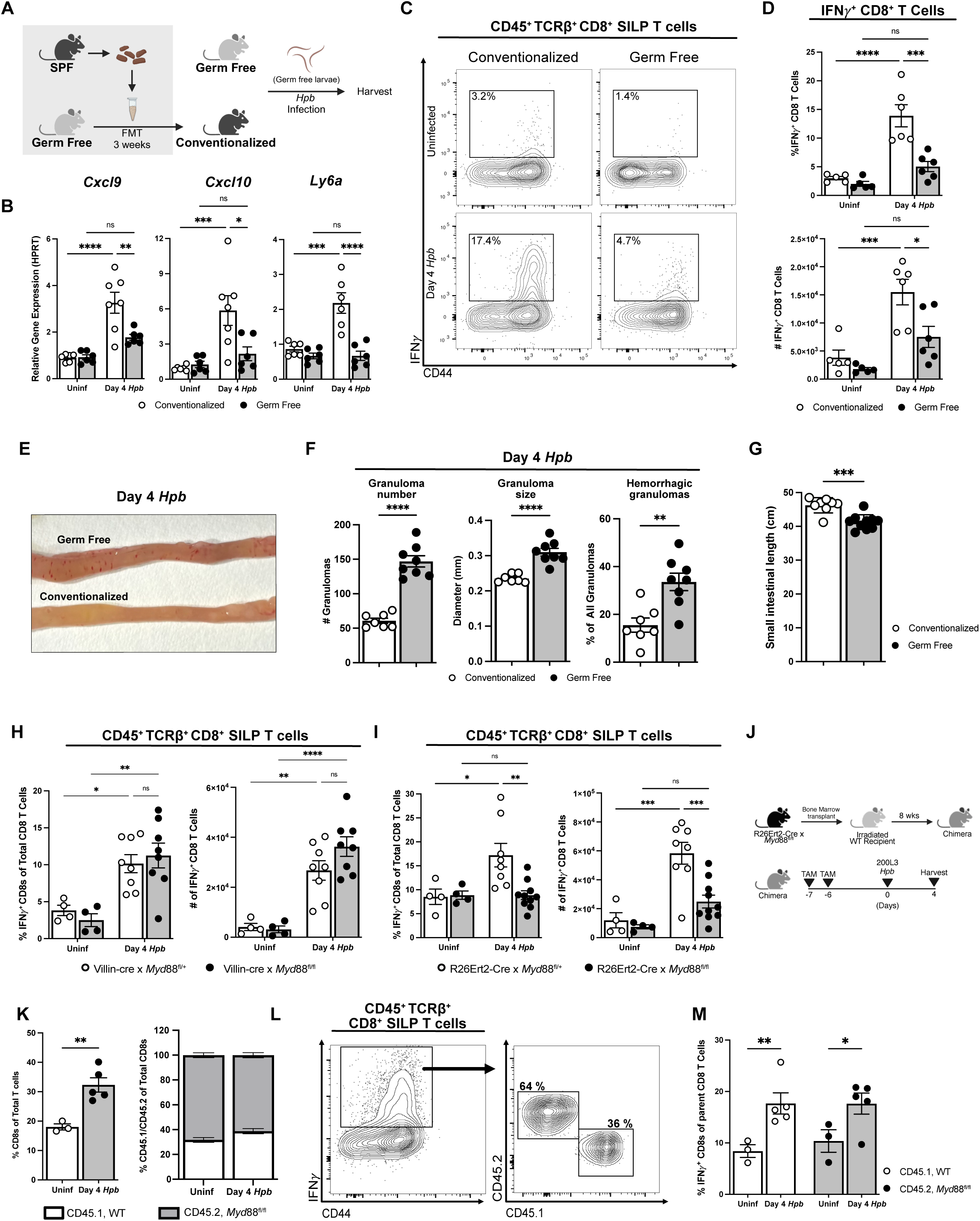
CD8^+^ T cell activation requires the gut microbiota and cell-extrinsic MyD88 signaling. (A) Schema of experimental approach for B-G. Germ-free (GF) mice were either gavaged with saline or fecal slurries from specific pathogen free (SPF) feces (i.e. conventionalized) three weeks prior to infection with 200 GF *Hpb* larvae. (B) ISG expression at day 4 *Hpb* infection in GF and conventionalized mice. (C) Representative flow cytometry plots from day 4 *Hpb* infected GF and conventionalized mice gated on viable CD45^+^ TCRβ^+^ CD8^+^ SILP cells, quantified in (D). Two independent pooled experiments, n=5-6 mice per group. (E) Representative image of day 4 *Hpb* infected conventionalized and GF intestines with granuloma number, size and frequency of hemorrhagic granulomas quantified in (F). (G) Quantification of small intestinal shortening of GF and conventionalized mice at day 8 *Hpb* infection with 600 larvae. (H-I) Deletion of MyD88 signaling in the (H) intestinal epithelium (*Villin*-cre x *Myd*88^fl/fl^) compared to heterozygous controls or (I) in whole body tamoxifen-inducible R26Ert2-cre x *Myd*88^fl/fl^ mice compared to heterozygous controls. In both experiments, SILP CD8^+^ T cell production of IFNγ was quantified following in vitro PMA/ionomycin stimulation at day 4 *Hpb* infection. Two independent pooled experiments, n=4-9 mice per group. (J) Schema for WT: R26Ert2-cre x *Myd*88^fl/fl^ (*Myd*88^-/-^) bone marrow chimeras. (K) Frequency of total CD8^+^ T cells in uninfected and day 4 *Hpb* infected mice (left) and the percentage host CD45.1 (*Myd*88^+/+^) and donor CD45.2 (*Myd*88^-/-^) cells from the total populations (right). (L) Representative flow cytometry plot of IFNγ^+^ CD8^+^ T cells, gated on viable CD45^+^ TCRβ^+^ CD8^+^ SILPs, and further gated into CD45.1^+^ and CD45.2^+^ cells. (M) Quantification of CD45.1 or CD45.2 IFNγ^+^ CD8 T cells from (M). Two independent pooled experiments, n=3-5 mice per group. Between group comparisons analyzed using student’s t test and time course analyses using 1- or 2-way ANOVA with Tukey’s posthoc analysis. *p<0.05, **p<0.01, ***p<0.001, and ****p>0.0001.

As a complementary approach, we examined the role of MyD88, an intracellular adaptor protein critical for downstream signaling of most toll-like receptor and IL-1R family members^17^. As a crucial cell interface between the microbiota and the immune system, we first deleted MyD88 signaling in the IEC compartment using Villin-cre x *Myd88*^fl/fl^ animals. We found epithelial-derived MyD88 expression to be dispensable for *Hpb*-induced IFNγ signaling (Figure 3H; S12A). By contrast, a systemic deletion approach using a tamoxifen-inducible Rosa26Ert2-cre x *Myd88*^fl/fl^ mice collectively demonstrated that MyD88 signaling was absolutely required for IFNγ signaling, in an epithelial cell-independent manner (Figure 3I; S12B). To test whether MyD88 signaling was intrinsic to the CD8^+^ T cell compartment, we generated irradiation chimeras in which bone marrow cells from tamoxifen-inducible Rosa26Ert2-cre x *Myd88*^fl/fl^ mice were transplanted into irradiated CD45.1 wildtype hosts (Figure 3J). Examination of these chimeras revealed an incomplete depletion of host CD8^+^ Trm cells residing in the SILP resulting in a mixed CD45.1 WT:CD45.2 *Myd88*^-/-^ hematopoietic compartment (Figure 3K). Examining these mice as mixed chimeras, we determined that the activation of CD8^+^ T cells occurred in a cell-extrinsic manner as IFNγ was produced by both CD45.1^+^ (WT-derived) and CD45.2^+^ (*Myd88*^-/-^-derived) cells (Figure 3L, M). Therefore, *Hpb*-induced tissue damage is not sufficient to induce IFNγ production, but rather depends on the presence of the gut microbiota and CD8^+^ Trm T cell-extrinsic MyD88 signaling.

### IFNγ signals recruit tissue-protective neutrophils to the site of *Hpb* infection

We next sought to understand how IFNγ signaling acts to limit tissue damage and promote disease tolerance to *Hpb* infection. Immunofluorescence imaging of *Hpb*-infected tissue showed CD8^+^ T cells in proximity to the granulomas (Figure 4A). Consistent with this observation, immunohistochemical analyses of day 2 *Hpb* infected tissue indicated activation of STAT1 (pSTAT), a transcription factor critical for IFNγ target gene expression, surrounding the granulomas in IFNγR competent but not receptor deficient mice (Figure 4B). Notably, pSTAT1 was detected in diverse cellular compartments including immune, stromal and epithelial cells. Since all cells express the IFNγR, we reanalyzed our previously published bulk RNAseq dataset from the intestine of day 2 *Hpb*-infected mice^6^ to survey which cell types or pathways may contribute to the tissue protective role of IFNγ production. Granulocyte migration, and neutrophil migration in particular, was highly upregulated as determined by gene ontology and gene set enrichment analysis (Figure 4C, D; S13). Highlighting genes associated with neutrophils and their associated chemokines, a volcano plot of all the differentially expressed genes at day 2 shows that *Cxcl5* was among the most highly upregulated (Figure 4E). Indeed, RNAScope analysis revealed that *Cxcl5* transcripts were enriched around the granuloma at day 4 in infected *Ifngr^+/+^* mice, but significantly decreased in *Ifngr*^-/-^ mice (Figure 4F). Consistently, we observed a striking accumulation of Ly6G^+^ neutrophils at the *Hpb* infection site that was abrogated in *Ifngr*^-/-^ mice (Figure 4G). Corroborating these results, neutrophil recruitment was also absent under GF conditions (Figure 4H), where IFNγ signaling was also eliminated (Figure 3D).

**Figure 4:**
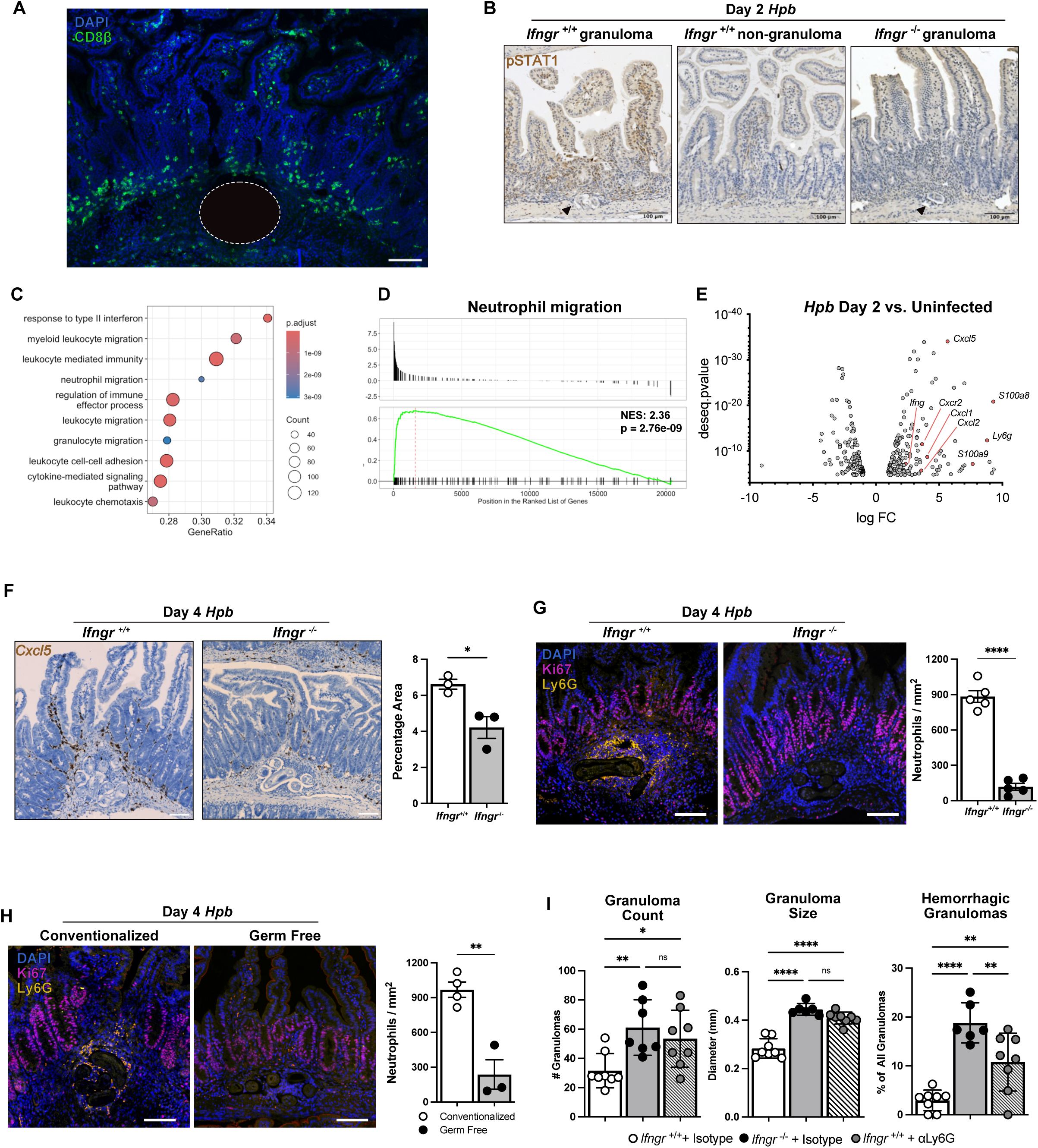
IFNγ-mediated neutrophil recruitment limits tissue damage to early *Hpb* infection. (A) Representative immunofluorescence image of CD8^+^ T cell localization in the granuloma region at day 4 *Hpb* in *Ifngr*^+/+^ mice. Granuloma is highlighted in dotted white circle. DAPI, blue; CD8β, green; scale bar, 100 µm. (B) Representative immunohistochemistry image of pSTAT1 staining in *Ifngr*^+/+^ granuloma (left), *Ifngr*^+/+^ non-granuloma (middle) and *Ifngr*^-/-^ mice (right) at day 2 *Hpb* infection. Black arrows indicate *Hpb* granuloma; scale bar, 100 µm. (C) Top differentially-expressed gene ontology terms at day 2 *Hpb* infection compared to uninfected controls following bulk RNA sequencing analysis of whole small intestinal tissue. Full analysis is provided in Figure S13. (D) GSEA analysis for neutrophil migration and (E) volcano plot highlighting upregulated neutrophil migration genes at day 2 *Hpb* infection compared to uninfected controls. FC, fold change. (F) *Cxcl5* RNAScope in granuloma areas of day 4 *Hpb* infected *Ifngr*^+/+^ and *Ifngr*^-/-^ mice, scale bar, 50 µm (left), quantified as the percentage of positive pixels per unit granuloma area of stain (right). Each dot is an average of 3-5 granulomas from three mice per genotype. Immunofluorescent imaging (Ly6G, yellow; Ki67, pink; DAPI, blue) of granulomas at day 4 *Hpb* infection in (G) *Ifngr*^+/+^ and *Ifngr*^-/-^ mice and (H) germ free and conventionalized mice. Scale bar, 50 µm. For each condition, two independent pooled experiments, 3-5 granulomas averaged per mouse with n=3-5 mice per group. (I) Number, size and frequency of hemorrhagic granulomas in isotype- or anti-Ly6G-treated mice from days -1 to +3 *Hpb* infection, two independent pooled experiments, n=7-10 mice per group. Between group comparisons analyzed with student’s t test. *p<0.05, **p<0.01, ***p<0.001 and ****p<0.0001.

Neutrophils are early responders to infection and are typically associated with aggravation of inflammation and microbicidal activity^18^. However, recent studies have indicated substantial heterogeneity in neutrophil populations^19^, including a growing appreciation for their tissue-protective functions^21,22^. To examine whether neutrophils impact tissue remodelling upon infiltration of the *Hpb* granuloma, we administered an anti-Ly6G neutrophil-depleting antibody from days -1 to 3 infection. Remarkably, neutrophil depletion recapitulated many features of infected germline *Ifngr*^-/-^ mice including an increase in the size, number and frequency of hemorrhagic granulomas (Figure 4I). We compared these results to *Ccr2*^-/-^ mice that lack Ly6C^+^ circulating inflammatory monocytes^23^. Importantly, *Ccr2*^-/-^ mice did not exhibit the same phenotype as neutrophil-depleted mice, consistent with their ability to retain IFNγ signaling and neutrophil recruitment (Figure S14A-C). However, neutrophil-depleted mice challenged with a high dose of *Hpb* larvae did not succumb to infection (Figure S14D). These results indicate that while IFNγ plays an important role in the recruitment of tissue-protective neutrophils, this cytokine must act on distinct cellular compartments to ensure host survival during high dose helminth infection.

### IFNγ signaling recruits neutrophils to helminth granulomas in a cell-extrinsic manner

Type 1 immunity is not typically associated with neutrophil recruitment. However, a recent study demonstrated that successful cancer immunotherapy was associated with a robust IFNγ gene signature in neutrophils that controlled tumor growth^22^. In this setting, anti-tumor neutrophils were identified as expressing low amounts of SiglecF compared to their pro-tumor counterparts^22^. To investigate whether this phenotype might also apply to neutrophils in the setting of infection, we first examined Sca-1 expression on infiltrating innate immune cells. Whereas Ly6C^+^ monocytes highly expressed Sca-1 in an IFNγR-dependent manner, neutrophils failed to express this IFNγ target gene (Figure 5A). Interestingly, when we assessed Siglec-F expression, neutrophils from *Ifngr^+/+^* mice expressed significantly less Siglec-F compared to the neutrophils we recovered from *Ifngr^-/-^* mice (Figure 5B) suggesting that they may be amenable to IFNγ-mediated immunomodulation. To definitively determine whether direct IFNγ signaling in neutrophils was required for their recruitment to *Hpb* granulomas, we constructed reciprocal irradiation chimeras where *Ifngr*^+/+^ or *Ifngr*^-/-^ bone marrow cells were transplanted into either irradiated *Ifngr*^+/+^ or *Ifngr*^-/-^ mice (WT-KO and KO-WT; Figure 5C). Genotype-matched (WT-WT and KO-KO) chimeras were generated as controls. As expected, there was a strong ISG signature at day 4 in the WT-WT control chimeras that was absent in the KO-KO group. Unexpectedly, the KO-WT group, with an IFNγR-competent radioresistant compartment, retained ISG expression while the WT-KO group did not (Figure 5D). Consistently, the loss of an IFNγ signature in the WT-KO chimeras, but not the KO-WT group, was associated with a defect in neutrophil recruitment (Figure 5E, F). These results indicated that neutrophils do not require direct IFNγ signaling for their recruitment. Further, it shows that direct IFNγ signaling on a radioresistant cell population is required for neutrophil recruitment.

**Figure 5:**
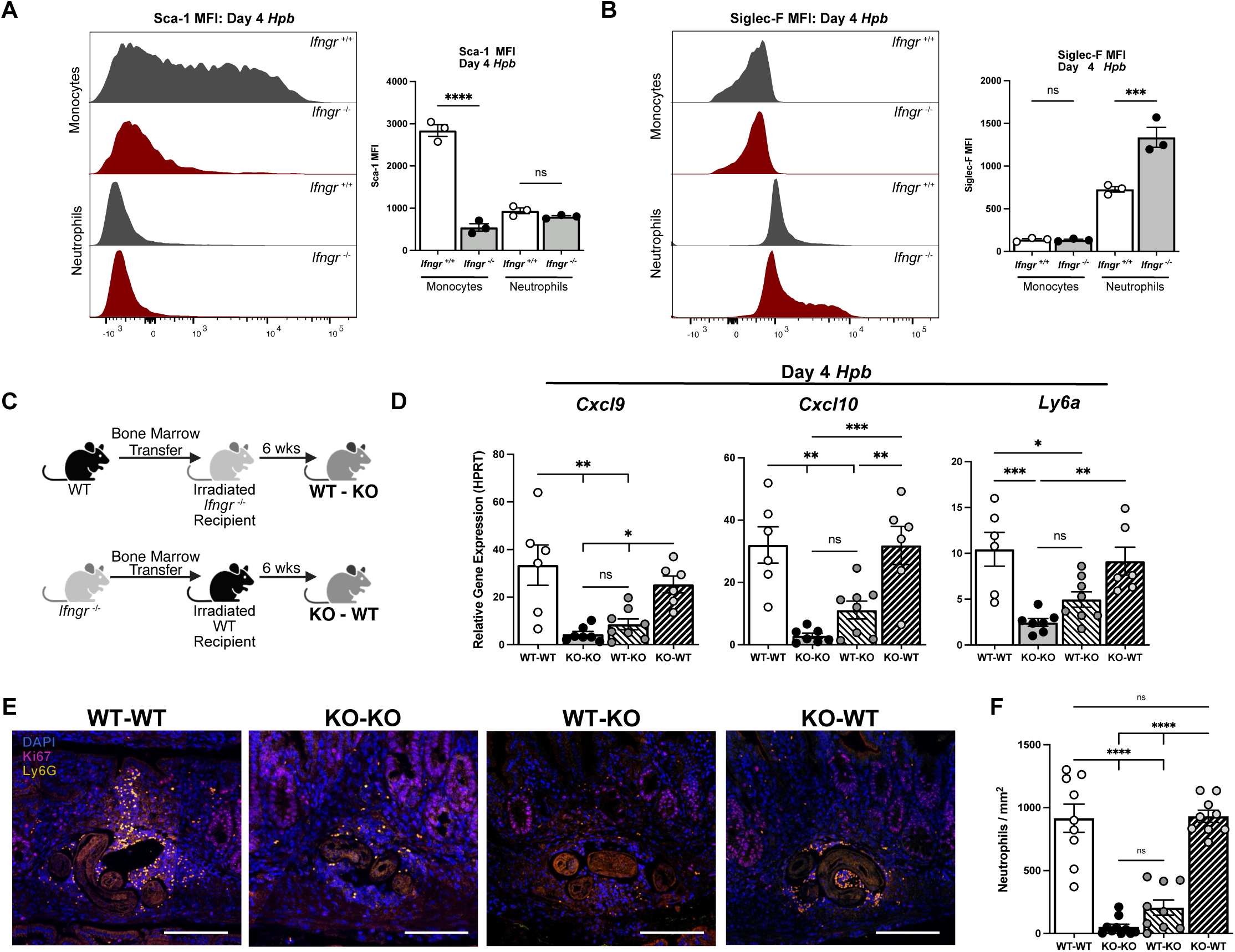
IFNγ recruits tissue-protective neutrophils to *Hpb* granulomas in a cell-extrinsic manner. (A, B) Representative histograms of (A) Sca-1 and (B) Siglec-F expression by monocytes (CD45^+^ CD11b^+^ MHCII^lo^ Ly6C^hi^) and neutrophils (CD45^+^ CD11b^+^ Ly6G^+^) at day 4 *Hpb* infection in *Ifngr*^+/+^ and *Ifngr*^-/-^ mice (left). Quantification of mean fluorescence intensity (MFI) of Sca-1 or Siglec-F expression by monocytes and neutrophils (right). (C) Schema of bone marrow chimera approach. (D) ISG expression, (E) representative immunofluorescence images (Ly6G^+^, yellow; Ki67, pink; DAPI, blue; scale bar, 50 µm) and (F) quantification of neutrophil recruitment in respective bone marrow chimeric groups. Two independent pooled experiments, n = 8-10 mice per group. Between group comparisons analyzed with student’s t test. *p<0.05, **p<0.01, ***p<0.001, and ****p<0.0001.

### Direct sensing of IFNγ by the intestinal stroma mediates neutrophil recruitment and tissue preservation during infection

Recently, the intestinal epithelium has been shown to integrate IFNγ signals to mediate protective effects and limit disease sequelae associated with bacterial infection and tumor immunity^24^. To test whether the intestinal epithelium was the radioresistant population sensitive to IFNγ signals during *Hpb* infection, we generated Villin-cre x *Ifngr*^fl/fl^ mice that failed to induce Sca-1 through IFNγ signaling in the intestinal epithelium (Figure S15A). At day 4, there were no defects in the early granulomatous damage phenotype (Figure S15B) nor was there elevated mortality in high dose infected mice compared to littermate controls (Figure S15C). Consistently, neutrophil recruitment was not compromised in Villin-cre x *Ifngr*^fl/fl^ mice (Figure S15D) indicating that the epithelium is not involved in IFNγ-mediated tissue protection or disease tolerance during *Hpb* infection.

An additional radioresistant population in mice are the stromal cells. The stroma is a heterogeneous cell compartment typically associated with production of the extracellular matrix^25^. Recently, the immunomodulatory properties of the stroma have become more appreciated in response to injury^26^, infection^27,28^ and in mesenchymal stem cell therapy^29^. As an initial examination of the intestinal stroma during *Hpb* infection, alpha smooth muscle actin (αSMA)-producing cells were visualized at day 4 by immunofluorescence microscopy (Figure 6A). As expected, the muscularis propria was densely populated by αSMA^+^ cells in *Ifngr*^+/+^ and *Ifngr*^-/-^ mice. However, in the absence of IFNγ signals, there was substantial remodelling of the stromal cells with significant accumulation of αSMA-producing cells surrounding the granuloma and extending throughout the intestinal villi that was not observed in *Hpb*-infected *Ifngr*^+/+^ mice. To determine whether direct IFNγ sensing by the stromal cell compartment regulated αSMA-producing cells, we crossed mice expressing a tamoxifen-inducible cre recombinase under control of the smooth muscle myosin Myh11 (Myh11Ert2-cre) with our *Ifngr*^fl/fl^ mouse line to generate Myh11Ert2-cre x *Ifngr*^fl/fl^ animals. Importantly, Myh11 and αSMA expression overlaps within the small intestinal mesenchyme^25^. Similar to the germline *Ifngr*^-/-^ mice, we observed an increase in αSMA staining in Myh11Ert2-cre x *Ifngr*^fl/fl^ but not Myh11Ert2-cre x *Ifngr*^fl/+^ littermate controls at day 4 (Figure 6B). Furthermore, eliminating IFNγ signaling in this stromal cell compartment reduced *Cxcl5* expression (Figure 6C) and neutrophil recruitment (Figure 6D) in the vicinity of the granuloma. Outcomes of parasite-induced damage that we observed in *Hpb*-infected germline *Ifngr*^-/-^ mice, GF mice and neutrophil-depleted mice including the number, size and frequency of hemorrhagic granulomas were all increased in Myh11Ert2-cre x *Ifngr*^fl/fl^ compared to littermate controls (Figure 6E). Similar results were observed in *Hpb*-infected mice lacking the IFNγR in a Collagen1A2-expressing stromal cells (Col1A2Ert2-cre x *Ifngr*^fl/fl^ mice; Figure S16). To assess susceptibility to a high dose challenge of *Hpb* larvae, elevated mortality was also observed in Myh11Ert2-cre x *Ifngr*^fl/fl^ compared to controls (Figure 6F), without any difference in worm burden between surviving animals (Figure 6G). Finally, assessment of intestinal transit and small intestinal length prior to the expected onset of mortality phenocopied the defects observed in germline *Ifngr*^-/-^ mice (Figure 6H-J). Collectively, our data indicate that direct IFNγ signaling into the intestinal stroma promotes disease tolerance to infection by recruiting tissue-protective neutrophils and restraining elaboration of the smooth muscle actin-expressing cell network to prevent pathological gut dysmotility.

**Figure 6:**
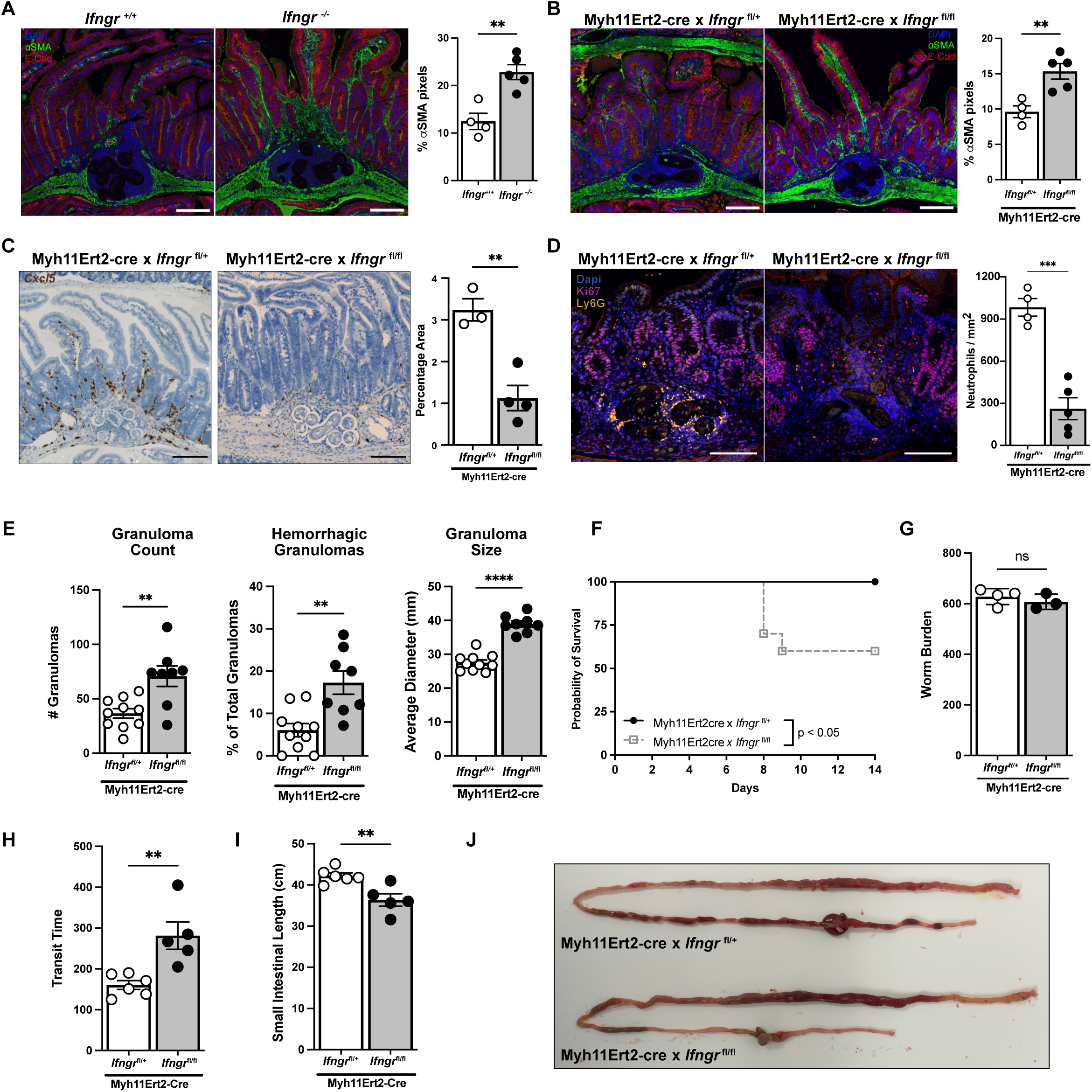
Stromal cell-intrinsic IFNγ signaling orchestrates disease tolerance to *Hpb* infection. (A) Representative immunofluorescence images of granuloma regions visualized with alpha smooth muscle actin (ɑSMA, green), E-cadherin (red) and DAPI (blue) staining (left) with quantification of the percentage of αSMA-positive pixels to total granuloma-associated tissue area (right). Scale bar, 100 µm; two independent pooled experiments with n=5-6 mice per group and each dot represented averaged 2-3 granulomas per mouse. (B) Same as (A) using Myh11Ert2-cre x *Ifngr*^fl/fl^ mice and controls (Myh11Ert2-cre x *Ifngr*^fl/+^). Two independent pooled experiments with n=5-6 mice per group and each dot represented averaged 2-3 granulomas per mouse. (C) *Cxcl5* RNAScope of Myh11Ert2-cre x *Ifngr*^fl/fl^ and controls at day 4 *Hpb* infection (left) with quantification of transcript-positive signal per unit area (right). Scale bar, 100 µm; One experiment, n=4-5 mice per group and each dot represented averaged 2-5 granulomas per mouse. (D) Immunofluorescent imaging (Ly6G, yellow; Ki67, pink; DAPI, blue; scale bar, 100 µm) in Myh11Ert2-cre x *Ifngr*^fl/fl^ mice or control animals (left) with quantification as in A (right). Two independent pooled experiments, n = 4-6 mice per group and each dot represented averaged 2-3 granulomas per mouse. (E) Quantification of visible granulomas as well as percentage of hemorrhagic granulomas and their size at day 4 *Hpb* infection between the indicated genotypes. Two independent pooled experiments, n=8-10 mice per group. (F) Kaplan-Meier survival curves of high dose (600 L3 larvae) Myh11Ert2-cre x *Ifngr*^fl/fl^ mice compared to controls with worm burden in surviving mice (G). Two independent pooled experiments, n=8-10 mice per group. Corresponding transit time (H), intestinal shortening (I) and a representative image of intestinal morphology (J). One experiment, n=5-6 per group. Between group comparisons analyzed with student’s t test. *p<0.05, **p<0.01, ***p<0.001, and ****p<0.0001.

## DISCUSSION

Type 1 immunity involves a multi-cellular sensing and signaling network indispensable for protection against intracellular pathogens^30^. A central effector molecule in this defense pathway is IFNγ, first identified by its potent ability to “interfere” with viral replication in leukocytes^31^. This function as well as many others, including the amplification of anti-bacterial and anti-protozoan immunity as well as a more recent measure of successful cancer immunotherapy, has continued to draw attention to this cytokine for over half a century^32^. However, pathogen elimination, also known as host resistance, is not the only defense strategy against infection. In fact, host resistance may involve an overreactive immune response leading to collateral damage of non-infected cells or tissues. Thus, a complementary or alternative defense strategy is disease tolerance. First described by plant biologists over a century ago^33^ and now being investigated in mammals including humans^34,35^, disease tolerance is defined as the host’s ability to control tissue damage without reducing pathogen load^1^. Although studies have identified immune cell-intrinsic checkpoints^36^ and metabolic resilience^37–39^ as ways to support this defense strategy, a role for immune-stromal cell networks has not been described. Our study reveals that, in addition to its robust anti-microbial activity, IFNγ is also fundamental to disease tolerance by preserving tissue function during tissue invasice infection. Specifically, we demonstrate that microbiota-dependent production of IFNγ by CD8^+^ Trm T cells acts directly on the intestinal stroma to recruit tissue-protective neutrophils and prevent pathological gut dysmotility during host invasion by an enteric helminth. Further, we demonstrate that these effects are largely independent of parasite burden, indicating that type 1 immunity mediates disease tolerance.

Despite the global prevalence of intestinal helminths, knowledge of the tissue response during the earliest events of infection remains nebulous compared to other pathogens. Here we use *Hpb* infection that follows a well-defined kinetic of infection limited to the small intestine to investigate how tissue invasion by a large, multicellular parasite is tolerated by the host. Our results build on previous studies, including our own, demonstrating that the early tissue-invasive phase of *Hpb* infection is defined by a type 1 immune response^6–8^. We now demonstrate that CD8^+^ αβ T cells residing in the lamina propria of the small intestine are the primary source of IFNγ production during early *Hpb* infection. Our results also indicate that, in this context, cognate antigen encounter was not required. However, the presence of the gut microbiota was critical, indicating that these cells were responding in an innate-like manner for host defense. Although antigen-independent differentiation and maintenance of CD8^+^ Trm T cells has been previously shown^14,40^, the ability of the microbiota to induce effector cytokine production by these cells to limit tissue damage has not been described, to our knowledge. While the specific microbes or microbial products remain to be identified, we found that hematopoietic cell sensing of the microbiota was required in a T cell-extrinsic, MyD88-dependent pathway. Additional gnotobiotic and cell-specific gene targeting approaches will need to be used to reveal the detailed signaling network leading to Trm T cell activation in this context.

Our studies also indicated a role for IFNγ in maintaining gut motility, a function not only important for host defense but also epithelial integrity. Intestinal dysfunction affects up to a third of the Western population with a significant proportion involving severe dysmotility^41^. Many of these conditions are associated recent intestinal infection, inflammatory events or post-surgical complications, while others remain ’idiopathic’^42^. Collectively, the basis for these intestinal dysfunction syndromes remains poorly understood. Upon high dose *Hpb* infection, we observed that mice with a germline or stromal cell-specific deletion of IFNγR demonstrated qualitative and quantitative small intestinal dysmotility. Interestingly, type 1 cytokines have been linked to hypocontractility of gut smooth muscle^43^ and IFNγ can directly inhibit αSMA production by arterial smooth muscle cells in vitro^44^. Functionally, IFNγ decreased longitudinal muscle contractility ex vivo^43^ and reduced contractibility and proliferation of human-derived intestinal smooth muscle cells^45^. Likewise, *Trichinella spiralis* infection of mice exhibiting elevated IFNγ production resulted in muscle hypocontractility^46^. The regulatory impact of IFNγ on smooth muscle contractility serves as an additional example of the mutual antagonistic nature of type 1 and type 2 cytokine responses^11^, as it is well-known that the latter including IL-4 and IL-13 enhance contractility to promote the weep and sweep response during helminth infection^47^. As such, it would be expected that the absence of IFNγR signaling would lead to increased helminth expulsion. However, we did not observe a decrease in worm burden at the time points examined, suggesting that induction of type 1 cytokine signaling is not simply used as an immune evasion strategy by the parasite but may be more relevant for limiting tissue damage. Consistent with these results, loss of IFNγR signaling in Sox10-expressing enteric glial cells during *Hpb* infection was recently shown to also compromise gut motility^8^. This study reported that enteric glial-intrinsic IFNγ signals promoted resolution of granulomas suggesting that IFNγ also contributes to tissue repair during chronic infection. Given the transient induction of IFNγ and its associated target genes during the early stage of infection, these data suggest that brief activation of type 1 immunity may have long-term impacts on wound healing, applicable to other settings of dysregulated repair and fibrosis such as ulcerative colitis.

Gut motility disorders are commonly associated with enteric nervous system (ENS) dysfunction and, in contrast to our results, CD8^+^ T cell production of IFNγ has been associated with aggravating dysmotility^48,49^. In these instances, the effects were due to direct regulation of enteric neuron density and signaling. However, distinct mechanisms underlie gut dysmotility that may or may not involve the ENS. Indeed, infection of mice with the roundworm *Nippostrongylus brasiliensis* led to hypercontractility of the small intestine that was independent of the ENS, confirming the importance of the stroma cell compartment as an additional mediator of gut motility^50^. Stratifying disease based on the form or cell types driving gut dysmotility may allow for more personalized approaches to effectively treating this morbidity.

We also observed a direct link between stroma-intrinsic IFNγR signaling and neutrophil recruitment to the site of infection. Stromal-neutrophil communication is an understudied field, though a battery of recent studies highlights the importance of the ECM in regulating leukocyte chemotaxis during mucosal tissue damage^51^. Our data indicates that IFNγ acts directly on the αSMA^+^ cells to mediate neutrophil chemotaxis through Cxcl5 expression. This stromal activity parallels a recent finding where following acute skin damage, an early responder subtype of stromal cells was identified that upregulated a set of immunomodulatory and repair transcripts^27^. Among these transcripts was the neutrophil chemokine *Cxcl5*, and stromal depletion of this cytokine early after injury lead to a prolongation of inflammation at later times^26^. Similarly, Cxcl12^+^ fibroblasts were shown to recruit neutrophils to the site of *Staphylococcus aureus* infection in skin through the detection of IL-17^27^. Additionally, successful cancer immunotherapy was recently associated with local neutrophil recruitment to the tumor site^22^ supporting the notion that IFNγ sensing by stromal cells could be a prime mediator of neutrophil recruitment.

Our data dovetail with studies showing a tissue-protective role for type 1 immunity in other contexts. For example, epithelial cell-intrinsic IFNγ signaling during intestinal bacterial infection and inflammation inhibits inflammasome activation responsible for driving pathogenic CD4^+^ T cell differentiation and colonic barrier damage^24^. Importantly, the tissue-protective effects of IFNγ had no impact on pathogen burden, indicating another setting in which IFNγ supports disease tolerance^24^. In addition, neutralization of IFNγ during pulmonary fungal infection of mice prevented lung damage and dysfunction while enhancing pathogen clearance, suggesting that IFNγ can simultaneously impact disease tolerance and host resistance pathways^52^. Moreover, IFNγ can also protect from immunopathology in non-infectious disease settings. Blockade of IFNγ signaling has been shown to limit alloreactive T cell responses during transplantation by inducing indolemine 2,3-dioxygenase in antigen presenting cells which blunts T cell expansion^53^. In addition, the presence of IFNγ is associated with neutrophil recruitment and regeneration following peripheral nerve injury^24,54^. Elegant in vitro studies have also revealed that human CD8^+^ T cell production of IFNγ can support epithelial tissue remodeling and wound healing by sensitizing cells to the tissue repair factor, amphiregulin^55^. Combining these studies with our data provide the impetus for re-evaluating how a cytokine, regarded to strictly drive inflammation and pathogen killing, can support adaptive tissue responses during injury and insult.

### Limitations of the study

Although our data identifies IFNγ signaling as critical for mitigating intestinal damage enabling disease tolerance to an enteric pathogen in the mouse model, whether these findings correlate to human chronic intestinal helminth infection and tolerance requires validation. Nonetheless, our findings identify a novel cellular network to guide further investigations. For example, evaluating the relevance of this network in other intestinal disorders of motility, such as the post-infectious state (’irritable bowel syndrome’)^56^ or the aged state^57^, may also shed light on these conditions. While our study demonstrates an important type 1 immune-stromal cell axis in disease tolerance by targeting smooth muscle myosin (Myh11)-expressing cells, the specific origins of this population within the broader stromal compartment remains to be determined. Additionally, investigation into helminth-derived molecules and their bioactivity on specific host cell types would provide a more holistic understanding of host-parasite interactions that might contribute to host health, pathogen persistence and disease tolerance.

## ACKNOWLEDGEMENTS

The authors would like to thank the staff in the Research Institute of the McGill University Health Centre Platforms including the Animal Resource Division and the Histopathological Core Facility. Special thanks to Cynthia Faubert and Catherine Hudon for their assistance with the germ-free animal studies and the McGill Interdisciplinary Initiative in Infection and Immunity for financial support to establish the Gnotobiotic Animal Research Platform at the RI-MUHC. S.W. is a recipient of a Canadian Institutes of Health Research (CIHR) Postdoctoral Fellowship (MFE-181759). I.L.K. is a recipient of a Fonds de Recherche du Québec Scholar Award. M.E.G. is a recipient of a Fonds de Recherche du Québec Postdoctoral Award. SW is currently funded by the Deutsche Forschungsgemeinschaft, DFG, project number WE 4971/6-1 and 9-1, the Excellence Cluster Balance of the Microverse (EXC 2051; 390713860), the Federal Ministry of Education and Research (BMBF) project number 01EN2001. This work was supported by the CIHR (PJT-178182) to I.L.K.

## AUTHOR CONTRIBUTIONS

Conceptualization, S.W., M.E.G., I.L.K.; Methodology, S.W., M.E.G., T.M.O., F.R., R.D.P., D.K.A., G.F., D.H., M.J.R., J.F.E.K., O.J.H., M.D., A.G., I.L.K.; Software, A.B.; Formal Analysis, S.W., M.E.G., T.M.O., A.B., D.K.A; I.L.K.; Investigation, S.W., M.E.G., T.M.O., F.R., R.D.P., D.K.A., G.M., G.F., I.L.K.; Resources, F.R., G.F., D.H., H.M., V.A., M.J.R., O.J.H., M.D., S.W., A.G., I.L.K.; Writing – Original Draft, S.W., I.L.K.; Writing – Review & Editing, All authors; Visualization, S.W., A.B., I.L.K.; Funding Acquisition, I.L.K.

## STAR METHODS

### RESOURCE AVAILABILITY

#### Lead Contact

Further information and requests for resources and reagents should be directed to and will be fulfilled by the lead contact, Irah King.

#### Materials availability

This study did not generate new unique reagents.

#### Data and code availability

- All the data has been previously published^6^. Microscopy data reported in this paper will be shared by the lead contact upon request.
- All original code is available in this paper’s supplemental information.
- Any additional information required to reanalyze the data reported in this paper is available from the lead contact upon request.

### EXPERIMENTAL MODEL AND STUDY PARTICIPANT DETAILS

#### Mice

All experimental procedures were performed in accordance with the McGill University Health Centre Research Institute Animal Resource Division with approved use protocol #7977. All mice were used between 6-10 weeks of age with an average weight of 18-25g. Wildtype (C57BL/6, No. 027) and CD45.1 (NCI B6-Ly5.1 No. 564) were obtained from Charles River. *Ifng*^-/-^ (No. 002287), GREAT (No. 017581), *Myd88*^fl/fl^ (No. 009108), *Ifngr* ^-/-^ mice (No. 003288) and *Ccr2* ^-/-^ mice (No. 017586), *Ifngr* ^fl/fl^ (No. 025394), *Villin*-cre (No. 004586), Myh11Ert2-cre (No. 019079), R26ERT2cre mice (No. 008463)and Col1A2Ert2-cre were all obtained from Jackson Labs. Germ free mice on a B6 background were purchased from the International Microbiome Centre at the University of Calgary. OT-I x Rag^-/-^ Ly5.1 were bred and maintained at our institution, TCRβKO mice were provided by Judith Mandl (McGill). IL-33-/-mice and CD4cre-IL36γ^fl/fl^ mice were bred and maintained at the University of Pennsylvania and Benaroya Institute, respectively. All mice were backcrossed to the C57BL/6 background and transgenic lines were bred in house. Mice were housed in groups (4-5 mice per cage) with *ad libidum* access to standard chow and water. Animals were housed at 22C with a 12h cycle of lights on (7am) and lights off (7pm). For all transgenic mice, littermates of the same sex were randomly assigned to experimental groups. For WT and germline transgenic models, mice of the same sex and age were randomly assigned to experimental groups. Sex had no effect on the experimental outputs, so all combined data is expressed as a combination of the sexes. Experiments involving germline gene-deficient mice were conducted at least once in adults co-housed (female mice) or given mixed bedding (male mice) for at least three weeks prior to experimentation to normalize microbiome effects. Uninfected and infected mice were not co-housed during an experiment. Health of the mice was monitored every 2-3 days or daily, in the case of high-dose infection studies. Any mouse that reached a body score of 2 or lost more than 20% of their initial weight was euthanized as a humane endpoint.

#### Cell lines

The mouse TLR4 reporter HEK293 cells (HEK-Blue mTLR4 cells) were engineered from the human embryonic kidney HEK293 cell line. Cells were used below 10 passages at 70-80% confluency. Source details can be found in the Key Resource Table.

#### Parasite

*Heligmosomoides polygyrus bakeri* is maintained in house.

## METHOD DETAILS

### Helminth infection and tamoxifen treatment

Infectious L3 larvae were propagated from SPF feces according to standard practices^58^. Unless indicated, mice were infected with 200 L3 larvae by gavage in sterile water. For the high-dose infection studies, mice were gavaged with 600 L3 larvae in sterile water. For inducible promoters, mice were gavaged with 5 mg of tamoxifen (Sigma Cat#T5648) dissolved in a 9:1 mixture of corn oil and ethanol. Treatments were delivered on day -7 and -6 days before infection.

### Antibody treatment

For the NK cell depletion experiments, mice were treated with i.p. 200 µg of anti-NK1.1 depletion antibody (clone PK136, BioXcell) or isotype (clone C1.18.4; BioXcell) on the day before infection and every second day until sacrifice^6^. FTY720 experiments were conducted by mice being treated i.p. with 1 µg/g of body weight of FTY720 in sterile water starting 4 days before infection and every other day until sacrifice. IL-18 was depleted using the anti-IL18 antibody (clone YIGIF74-1G7), at a dose of i.p. 200 µg per mouse starting 1 day before infection and every other day until sacrifice. IL-15 was similarly neutralized using i.p. 200 µg of the M96 clone with the IgG2a (clone C1.18.4) as a control.

### Small molecular treatment experiments

TRITC-Dextran (Sigma Cat#T1037) was administered i.v. at 50 mg/kg in sterile physiological saline 10 min before sacrifice. Tissue for histology was prepared as previously described for immunofluorescence. FITC-Dextran (Sigma Cat#FD4) was administered to 3 h fasted mice at a dose of 12 mg in 150 µl of PBS by oral gavage. Food was replaced and mice were sacrificed 4 h later by cardiac puncture. Plasma was isolated and FITC signal was read immediately on a Tecan infinite M200 plate reader. Dosing of rIFNγ was conducted by i.p. treating mice with 1µg of rIFNγ (Peprotech, Cat#315-05) daily from the day of infection until the day of sacrifice.

### Adoptive transfer experiments

All adoptive transfer experiments followed the same protocol. Donor cells were isolated from the organ indicated and either the complete cell compartment or an isolated cell population was taken. For the adoptive transfer of WT and *Ifng*^-/-^ T cells into TCRβ^-/-^ mice, T cells from the spleen and LNs were isolated using Stem cell EasySep Mouse T Cell isolation kit (Cat#19851) and a total of 5 x 10^6^ cells were transferred to recipient TCRβKO mice. For the adoptive transfer of OT-1xRag1^-/-^ cells into Rag1^-/-^ mice, whole spleen from OT-1 Rag1^-/-^ mice was isolated and 5 million cells were transferred to CD45.2 Rag1^-/-^ mice. For the controls, CD8 T cells were isolated from the spleens of CD45.2 WT mice using the EasySep Stem Cell Kit for CD8 T cells (Cat#19853C.1) and 5 million cells were transferred into Rag1^-/-^ mice. P14 cell adoptive transfer and LCMV infection was conducted as previously described^59^. Mice were allowed to acclimatize for 6 weeks to the new cell population before experiments were conducted.

### Bone marrow chimeras

Recipient mice were irradiated (X-RAD SmART Irradiator) twice, 4 h apart with 5.5 Gy of irradiation for a final lethal dose of 11 Gy. Immediately after the second dose, mice were i.v. injected with total bone marrow cells from donor mice with 5 million cells/mouse. Mice were allowed to reach chimerism for 8 weeks before infection with *Hpb*.

### Germ free mice experiments

Fecal microbiota transfers (FMT) were performed by taking SPF feces from sex-matched adult WT mice and diluting 160 times in sterile reduced PBS. The slurry was filtered through 100 µm mesh filters and 250 µL was gavaged into germ free (GF) mice. Mice were allowed to conventionalize for 3 weeks before infection. GF larvae were prepared from adult SPF mice as previously described (Russel 2020, Star Methods). Both GF and conventionalized mice were infected with sterile *Hpb* larvae as described above.

### Granuloma characterization

The top 5-10 cm of the proximal duodenum was harvested and placed in sterile HBSS supplemented with 5 % FBS. Intestines were cut longitudinally and under a Micromaster dissection microscope (Fisher scientific), visible granulomas were counted and scored for their hemorrhagic nature. The granuloma diameter was measured using an electric caliper (Electronix Digital Caliper, IP54) and each point is representative of the average size of 20-30 granulomas found within the tissue processed.

### Carmine red assay

Unfasted mice were orally gavaged with 200µL of 6% carmine red, prepared in PBS. Fecal pellets were monitored continuously until the first pellet with a full red color was passed by each mouse while the time for the first pellet to emerge was considered to be transit time.

### Cell Extraction and flow cytometry

The proximal 5-10cm of the duodenum was used to extract lamina propria (LP) or intestinal epithelial cells (IECs) as previously described^6^. In brief, fat and peyer’s patches were removed from the tissue and subsequently washed in Hank’s Balanced Salt Solution (HBSS) before two 20 min incubations at 37°C in 5mM EDTA buffer in HBSS supplemented with 10% heat inactivated fetal bovine serum (FBS) and 15mM HEPES. The tissue was filtered through a 100 µm filter to collect the IEC compartment. The remaining tissue was wash twice in HBSS with 10% FBS and digested in RPMI 1640 supplemented with 10% FBS, 15mM HEPES, 100U/mL of DNase and 200 U/mL of collagenase VIII for 23 min at 37°C with shaking at 250rpm. The digestion was stopped by adding 35 mL of cold R10 buffer (RPMI 1640 supplemented with 10% FBS, 15mM HEPES, 1% L-glutamine and 1% penicillin/streptomycin). The tissue was manually crushed, passed through a 100 µm filter and centrifuged at 1800 rpm for 8 min at 4°C and resuspended in R10 buffer. For experiments where cells were stimulated, 3 million cells were washed twice in sterile PBS µ resuspending them in PMA (100 nM), ionomycin (1 µm) and GolgiStop (0.67uL/mL) for 4h at 37°C. The cell suspension was incubated with a fixable viability dye (eFluor 506) for 20 min at 4°C. Cells were washed and incubated with Fc block (10 min at 4°C) following be surface staining (30 min at 4°C). Cells were fixed and permeabilized with the FoxP3 Fix/Perm kit for 30 min at 4°C according to manufacturer’s instructions and intercellular staining was performed in the kit’s permeabilization buffer (45 min at 4°C). Data acquisition was performed with a LSR Fortessa (BD Biosciences) and analyzed using FlowJo software (BD Biosciences).

### Immunofluorescence

Freshly dissected tissues were cut longitudinally, washed in cold PBS and immediately prepared into a “Swiss roll” and placed in a histological cassette. The tissue was fixed in10 % formalin solution for 24 h before being rinsed and transferred to 70 % ethanol. Sections were paraffin embedded by the histological core at the RI-MUHC and cut into 5 µm section using a microtome (Leica). Sections were deparaffinized in sequential xylenes followed by serial hydration steps in descending concentrations of ethanol. Antigen retrieval was done in citrate buffer for 20 min in a steamer followed by rest in then hot citrate buffer for 20 min at room temperature. Slides were washed 3 times for 5 min in PBS before blocking with 3 % bovine serum albumin (BSA) in PBS for 3 h. Samples were left with the primary antibody (indicated in key resource table) for 24 h at 4 °C followed by a secondary antibody stain (indicated in key resource table) for 2 h at room temperature in a hydration chamber. Slides were mounted in Prolong diamond antifade mountant (Invitrogen, Cat#P36970) and allowed to cure at room temperature overnight before imaging on Zeiss confocal microscope (LSM700). Images analysis was done in ImageJ (NIH) and in QuPath (Bankhead et al., 2017). Enumeration of neutrophils within the granuloma was done manually and represented as the number of neutrophils per granuloma area. ɑSMA quantification was conducted in ImageJ (NIH) was quantifying the number of positive ɑSMA pixels per granuloma associated region, including the affiliated villi, and normalizing that to the total area of granuloma associated tissue. Quantification of the number of goblet and tuft cells was determined manually, compared to an automatically selected outline of villi regions in the granuloma associated regions using an automatic function in Qupath.

### Immunohistochemistry

Slides were prepared as described for immunofluorescence. Slides were deparaffinized and rehydrated in several gradients of descending ethanol. Antigen retrieval was performed for 20 min at >90°C in pre-warmed citrate buffer in a steamer. Samples were allowed to rest in the citrate buffer for 20 min outside of the steamer followed by quenching in 3 % hydrogen peroxide for 15 min and one 10 min wash in PBS. Samples were blocked in 1 % BSA in PBS for 3 h in a hydration chamber followed by primary antibody staining with pStat1 (Tyr701, Cell Signaling #9167) at 1:200 dilution overnight at 4 °C. After 3, 20 min washes in PBS, a secondary antibody stain with a ready-mix rabbit HRP antibody (Dako Cat#K4003) was performed for 90 min at room temperature. After 3, 20 min washes in PBS, staining was revealed with 3,3’-Diaminobenzidine (DAB) reagent (Sigma Cat#D3939) according to manufacturer’s instructions. Images were further counterstained with hemoxylin, followed by bluing in acidified water and dehydration and clearing in xylene. Slides were mounted in a xylene-based mountant (Cytoseal; Thermofisher) and imaged on an Axio Imager M2 microscope (Zeiss) with 20X magnification.

### RNAScope

RNAScope was performed according to manufacturer instructions (ACD bio, Cat#322310). The RNAscope probe *Cxcl5* (Cat#467441) was used. Images were acquired with an Axio Imager M2 microscope (Zeiss) at 20X magnification. Images were quantified using the pixel count function of Qupath, a digital pathology image analysis software^60^.

### RNA extraction and qPCR

RNA was extracted from 0.5 cm of tissue using Tri reagent (Sigma-Aldrich) following manufacturer’s instructions, with an additional overnight ethanol precipitation step. Briefly, cleaned RNA was resuspended in 90 µl of water with 10 µl of 3M ammonium acetate and combined with 300 µl of ice cold ethanol and left at -20°C for 18 h. Purified RNA was centrifuged, washed in 70% ethanol and resuspended in a final volume of RNase/DNase-free water. Reverse transcript was performed with AdvanTech 5x Reverse Transcription Mastermix (Cat# AD100-31404) and quantitative PCR was conducted using Advance Tech QPCR Mix (Advantech, Cat# AD100-21 402) and analyzed relative to HPRT internal controls using the 2^-ΔΔCT method. All primer pairs are listed in the key resources table.

### Ex vivo stimulation

Small intestinal cell preparations were made according to the protocol for FACS. Single cells were placed in a 24 well plate at 3 x 10^6^ cells in T cell media containing RPMI, 10% FBS, penicillin/streptomycin and 5mM HEPES. Cells were stimulated with a combination of IL-2 (0.55 ng/mL), IL-12 (10 ng/mL), IL-33 (25 ng/mL), IL-18 (10 ng/mL) and/or IL-36γ (10 ng/mL) for 18 h at 37 °C). 4 h before the end of the stimulation, GolgiStop was added to the cells.

### Tissue Bacterial Burden

Systemic bacterial burden was determined by taking whole tissue slurries of the lung, liver and spleen prepared by manually crushing the tissue through a 100 µm filter. The suspension was made up to 5 ml and 100 µl of this solution, or its dilutions, was plated on TSA plates with 5 % defibrillated sheep’s blood. The TSA base was composed of 10 g of Peptone, 10 g of yeast, 5 g of NaCl and 15 g agar per liter of water. Plates were incubated aerobically for 48 h before enumeration of colonies manually.

### Serology

Plasma was stored at -80°C until further analysis. Measurements of conventional serological parameters were performed by SYNLAB.vet GmbH (Leipzig, Germany).

### Cytokine measurements

Plasma cytokines were determined in duplicates using the LEGENDplex™ Mouse Inflammation Panel 13-plex (BioLegend, San Diego) on a BD Accuri™ C6 Plus Flow Cytometer (BD Biosciences) according to manufacturer instructions.

## QUANTIFICATION AND STATISTICAL ANALYSIS

### Bulk RNA seq analysis

Reanalysis of the RNASeq dataset was conducted in R using ClusterProfiler and DOSE R packages. Code is available in the supplementary information.

### Statistical analysis

Details of the statistical analyses in terms of group sizes and specific tests used can be found in the figure legends. All data are represented at the mean results derived from independent mice with the spread of data represented as the standard error of the mean (SEM) with significance determined when p < 0.05. GraphPad Prism 10 software was used to perform statistical analyses. Student’s t test, one- or two-way ANOVA were used as appropriate. Nonparametric tests were used when the results did not fit a normal distribution.

## SUPPLEMENTAL INFORMATION

### Supplemental Figure Legends

**Figure S1: The type 2 immune signature is enhanced in *Ifngr*^-/-^ mice during *Hpb* infection.**

(A) Tuft and goblet cells were enumerated in granuloma-associated regions of *Ifngr*^+/+^ and *Ifngr*^-/-^ mice at 6 and 8 days *Hpb* infection by immunofluorescence of DCLK and Lectin labeling, respectively. Quantification represents the average number of tuft or goblet cells per unit area of the crypt/villus axis surrounding the granuloma. Two pooled independent samples, n = 5-8 mice. (B) Representative flow cytometry plots of Gata3^+^ CD4 Th2 cells at day 6 *Hpb* infection (left), with quantification across multiple mice (right). Gated on live CD45^+^TCRβ^+^CD4^+^ SILP cells. Representative plot of two independent experiments, n = 4-5 mice per group. Time course analyses were conducted with 2-way ANOVA and Tukey’s posthoc analyses. *p<0.05, **p<0.01, ***p<0.001.

**Figure S2: IFNγ differentially affects intestinal barrier integrity during *Hpb* infection.**

Intravenously-administered TRITC-Dextran dye was administered to 4 day *Hpb* infected mice 10 min before sacrifice. (A) Representative immunofluorescent images of *Ifngr*^+/+^ and *Ifngr*^-/-^ mice showing TRITC infiltration (yellow) into the granuloma (left) with quantification (right). Quantification represents fraction of fluorescent area with a TRITC-positive signal normalized to the total area of the granulomatous area. Scale bar, 100 µm; One experiment, n = 3 mice per group. (B) Granuloma damage characterization in the small intestine following low-dose recombinant IFNγ treatment of C57BL/6 mice until the time of sacrifice. Two pooled independent experiments, n = 6 mice per group. (C) Small intestinal barrier integrity was tested by gavaging mice with FITC-Dextran molecules and measuring FITC signal in the serum, 4 h after administration. One experiment, n = 3 mice per group. (D) Indirect endotoxin measurement from *Hpb-*infected mice detected from plasma with a TLR4-HEK-Blue cell reporter line. Semi-quantitative data is represented as the OD 655 from the cell supernatant, 18 h after plasma exposure. Two independent pooled experiments, n = 7 mice per group. Between group comparisons analyzed with student’s t test and with time course analyses with 2-way ANOVA and Tukey’s posthoc analysis. *p<0.05, **p<0.01, ***p<0.001.

**Figure S3: Systemic response to high-dose *Hpb* infection**

(A) Percent weight change over time of *Ifngr*^+/+^ and *Ifngr*^-/-^ mice following 600 *Hpb* larvae challenge. Two independent pooled experiments, n = 15 mice per group. (B) Total culturable bacterial burden from high-dose infected mice at day 8 post *Hpb* infection in the whole spleen, liver and lung. Three pooled independent experiments, n = 10-14 mice per group. (C) Serum serology report of high dose *Hpb* infected mice at day 8. (D) Cytokine profile of the same groups shown in C. (E) Circulating endotoxin levels in high-dose *Hpb* infected mice measured semi-quantitatively by TLR4-HEK-Blue reporter cells. Serology and cytokines conducted using n = 9-10 mice per group. ALT: alanine transaminase, GLDH: glutamate dehydrogenase, LDH: lactate dehydrogenase, CK: creatine kinase. Between group comparisons analyzed with student’s t. *p<0.05, **p<0.01, ***p<0.001.

**Figure S4: IFNγ production from SILP CD8^+^ T cells following in vitro stimulation**

(A) Representative flow cytometry plots of SILP IFNγ^+^ CD8 T cells in uninfected or days 2 or 4 post *Hpb* infection WT mice. Gated on live, CD45^+^ TCRβ^+^ CD8^+^ T cells. (B, C) Frequency and number of IFNγ^+^ of SILP (B) CD8^+^ and CD4^+^ (C) T cells from the indicated time point of *Hpb* infection. (D) Frequency and number of of Sca1^+^ EpCAM^+^ epithelial cells as determined by flow cytometry at the indicated time point of infection. Two independent experiments, n = 4-6 per group. Time course analyses were conducted with 1-way ANOVA and Tukey’s posthoc analysis. *p<0.05, **p<0.01, ***p<0.001.

**Figure S5: Immunophenotyping of IFNγ^+^ CD8^+^ T Cells**

eYFP^+^ and eYFP^-^ CD8^+^ T cells were gated from live, CD45^+^ TCRβ^+^ cells at day 2 *Hpb* infection and the mean fluorescence intensity (MFI) of a selection of T cell activation and tissue-resident markers was assessed. Two independent pooled experiments, n = 5 mice per group. Between group comparisons analyzed with student’s t. *p<0.05, **p<0.01, ***p<0.001.

**Figure S6: SILP cell preparation contain few vascular CD8^+^ T cells**

(A) Representative flow cytometry plot of viable T cells found in the SILP, spleen or circulating blood preparation based on in vivo CD45.2 intravascular labeling. Gated on live, total CD45^+^ TCRβ^+^ T Cells. (B) Quantification of intravascular labelled (CD45.2+ for both fluorophores) CD8^+^ T cells within the SILP and (C) spleen represented as the frequency of intravascularly labelled cells within the total CD8^+^ T cell population. Two independent pooled experiments, n = 6 per group. Time course analyses were conducted with 1-way ANOVA and Tukey’s posthoc analysis. *p<0.05, **p<0.01, ***p<0.001.

**Figure S7: Multi-tissue detection of IFNγ-expressing (eYFP^+^) CD8^+^ T cells during *Hpb* infection**

At days 2 and 4 *Hpb* infection, the indicated tissues were harvested for detection of IFNγ-expressing (eYFP^+^) CD8^+^ T cells. In all cases, samples are gated on live, CD45^+^ TCRβ^+^ CD8^+^ T cells and frequencies are represented as eYFP^+^ cells as a percentage of total CD8^+^ T cells. IELs: intestinal epithelial lymphocytes, mLNs: mesenteric lymph nodes, BM: bone marrow, PP: Peyer’s patches. Two independent pooled experiments, n = 6 mice per group. Time course analyses were conducted with 1-way ANOVA and Tukey’s posthoc analysis. *p<0.05, **p<0.01, ***p<0.001.

**Figure S8: SILP accumulation of ɑ4β7^+^ CD8^+^ T cells occurs after initiation of IFNγ target gene expression.**

(A) Representative flow cytometry plots of CD44^+^a4β7^+^ CD8^+^ T cells at the indicated time point of *Hpb* infection. Gated on live, CD45^+^ TCRβ^+^ CD8^+^ SILP T Cells. (B) Quantification of the frequency and number of ɑ4β7^+^ CD8 T cells in the SILP. One experiment, n = 3 mice per group. Time course analyses were conducted with 1-way ANOVA and Tukey’s posthoc analysis. **p<0.01, ***p<0.001.

**Figure S9: LCMV-specific memory CD8^+^ T cells are activated independent of cognate antigen during *Hpb* infection**

(A) Schema of experimental approach. Purified LCMV-specific Thy1.1^+^ P14 cells were adoptively transferred into Thy1.1^-^ WT mice. Mice were infected with LCMV Armstrong. 30 days later, recipient mice were infected with *Hpb*. (B) Representative flow cytometry plots of uninfected and day 4 *Hpb* infected mice, gated on live, CD45^+^ TCRβ^+^ CD8^+^ SILP T Cells. (C) Quantification of Ki67^+^ Thy1.1^-^ (recipient) and Thy1.1^+^ (donor P14) cells following *Hpb* infection at day 4. Two independent pooled experiments, n = 4-8 per group. Between group comparisons analyzed with student’s t. *p<0.05, **p<0.01, ***p<0.001.

**Figure S10: IL-18, IL-33 and IL-36y signaling are dispensable for IFNγ production by CD8^+^ T cells following *Hpb* infection**

(A) qPCR analysis of *Il18, Il33* and *Il36g* mRNA expression during early *Hpb* infection. Three independent pooled experiments, n = 5-6 per group. (B, C) In vitro stimulation of uninfected and day 2 *Hpb* infected SILP cells for 18h with the indicated combination of cytokines. Two independent experiments, n = 4 mice per group. (D) Gene expression analysis of day 2 *Hpb* infected tissue from WT and IL-33^-/-^ mice. One independent experiment, n = 4 mice per group. (E) IFNγ production by CD8^+^ T cells from CD4-cre x *Il36g*^fl/fl^ *Hpb* infected mice. (F) Frequency and cell count of eYFP^+^ CD8^+^ T cells at day 2 and day 4 *Hpb* infection, following administration of neutralizing anti-IL-18 (nIL18) antibody administration. Two independent experiments, n = 3-5 mice per group. Between group comparisons analyzed with student’s t test and with time course analyses with 1-way ANOVA and Tukey’s posthoc analysis. *p<0.05, **p<0.01, ***p<0.001.

**Figure S11: IL-15 is dispensable for IFNγ production by CD8^+^ T cells following *Hpb* infection IFNγ-**reporter mice were administered an anti-IL-15 neutralizing antibody prior to *Hpb* infection. It’s bio-activity was verified by the complete depletion of NK cells in the (A) blood and (B) SILP. (C) Quantification of eYFP^+^ CD8^+^ T cells in the SILP following treatment. Gated on live, CD45^+^ TCRβ^+^ eYFP^+^ CD8^+^ T cells (left). Quantification of Sca1^+^ epithelial cells in the IEC fraction. Gated on live, CD45-EpCAM^+^ epithelial cells (right). Two pooled independent experiments, n = 7-8 per group. Between group comparisons analyzed with student’s t test. *p<0.05, **p<0.01, ***p<0.001.

Figure S12: MyD88 signaling in the intestinal epithelium is dispensable for IFNγ target gene expression following *Hpb* infection

(A) ISG expression in Villin-cre x MyD88^fl/fl^ mice at days 2 and 4 post *Hpb* infection compared to controls. (B) ISG expression in R26Ert2cre x MyD88^fl/fl^ mice at days 2 and 4 post *Hpb* infection compared to controls. Two independent pooled experiments were performed for each genotype, n = 8 mice per group. Time course analyses conducted with 2-way ANOVA and Tukey’s posthoc analysis. *p<0.05, **p<0.01, ***p<0.001.

**Figure S13: Unmodified GSEA analysis from day 2 *Hpb* infected mice**

Top 20 differentially expressed GO pathways at day 2 *Hpb* infection compared to uninfected controls.

**Figure S14: Deletion of CCR2 does not impact IFN**γ **production, neutrophil recruitment or *Hpb* granuloma-associated tissue remodeling.**

(A) In vitro IFNγ production by CD8^+^ T cells following *Hpb* infection isolated from the SILP of *Ccr2*^-/-^ mice and controls. Two independent pooled experiments, n = 4-8 per group. (B) Representative immunofluorescence images of neutrophil accumulation to the site of granulomas in *Ccr2*^-/-^ mice, and their controls, at day 4 post *Hpb* infection. Ly6G, yellow; Ki67, pink; and DAPI, blue; scale bar, 200 µm. (C) Characterization of granuloma-associated tissue remodeling in *Ccr2*^-/-^ and control mice at day 4 *Hpb* infection. Two independent pooled experiments, n = 6 mice per group. (D) Survival plot (left) and worm burden (right) of high dose infected mice treated with anti-Ly6G antibody from days -1 to 5 of *Hpb* infection. Two independent pooled experiments, n = 8-10 per group. Between group analyses conducted with student’s t-test while time course analyses conducted with 2-way ANOVA and Tukey’s posthoc analysis. *p<0.05, **p<0.01, ***p<0.001.

**Figure S15: Epithelial IFNγ signaling is dispensable for disease tolerance to *Hpb* infection**

(A) Sca-1 expression by intestinal epithelial cells (IECs) in Villin-cre x *Ifngr*^fl/fl^ mice and control animals at days 2 and 4 *Hpb* infection. Gated on live, CD45^-^ EpCAM^+^ cells. Two independent pooled experiments, n = 4-6 per group. (B) Enumeration of granuloma counts, size and hemorrhaging Villin-cre x *Ifngr*^fl/fl^ mice and control animals at day 4 *Hpb* infection. Two independent pooled experiments, n = 5-6 mice per group. (C) Survival plot of high dose infected Villin-cre x *Ifngr*^fl/fl^ mice and control animals. (D) Representative immunofluorescence imaging (left) and quantification (right) of neutrophil recruitment in Villin-cre x *Ifngr*^fl/fl^ mice and control animals at day 4 post *Hpb* infection. Two independent pooled experiments, n = 6 mice per group. Between group analyses conducted with student’s t-test while time course analyses conducted with 2-way ANOVA and Tukey’s posthoc analysis. *p<0.05, **p<0.01, ***p<0.001.

**Figure S16: IFNγ signaling in Col1A2-expressing intestinal cells restricts *Hpb*-induced granuloma-associated tissue remodeling.** Quantification of granuloma counts, size and hemorrhaging in Col1A2Ert2-cre x *Ifngr*^fl/fl^ and Col1A2Ert2-cre x *Ifngr*^fl/+^ mice at day 4 *Hpb* infection. n = 4-6 mice per group. Between group analyses conducted with student’s t-test. *p<0.05, **p<0.01, ***p<0.001.

## GSEA analysis raw code

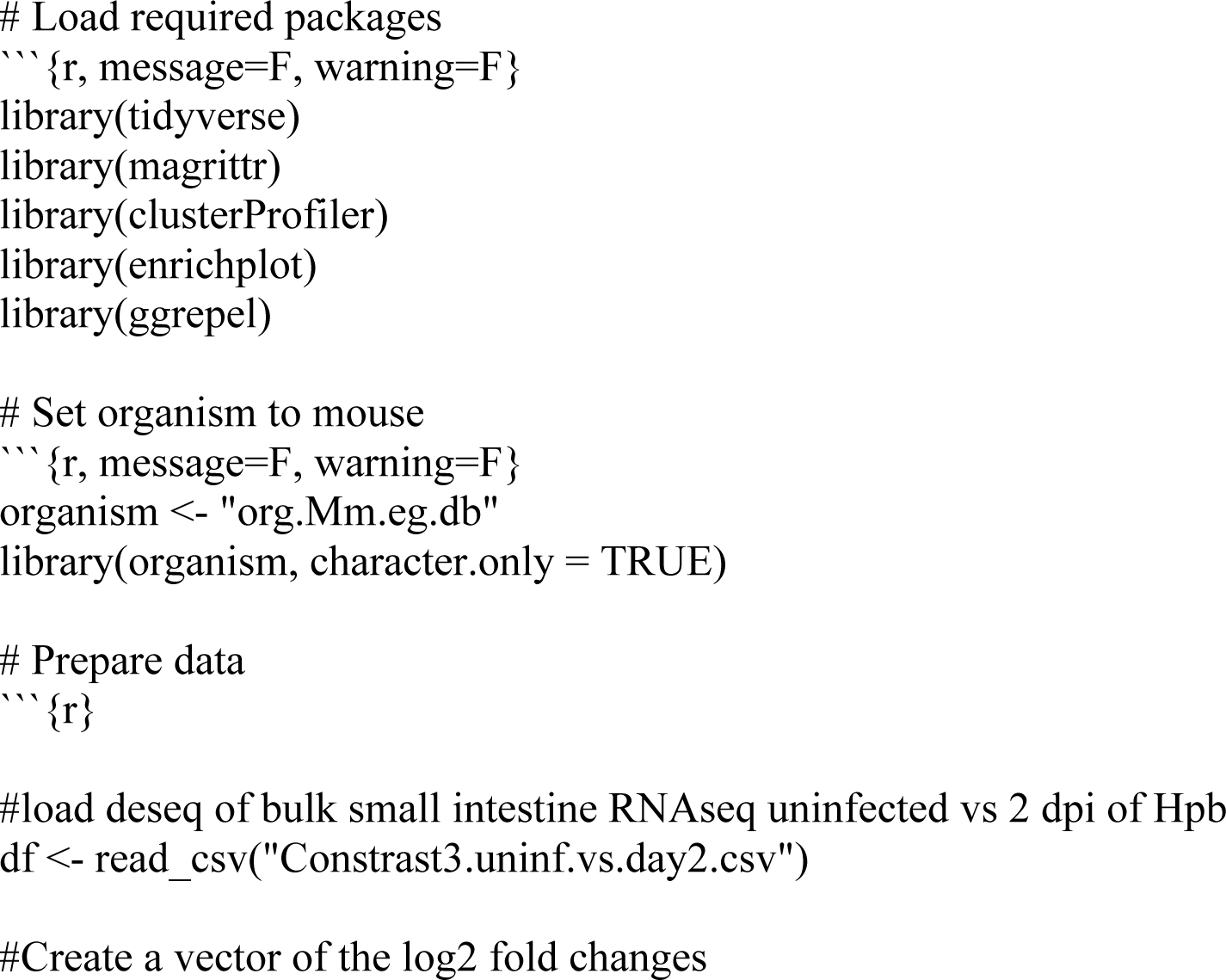
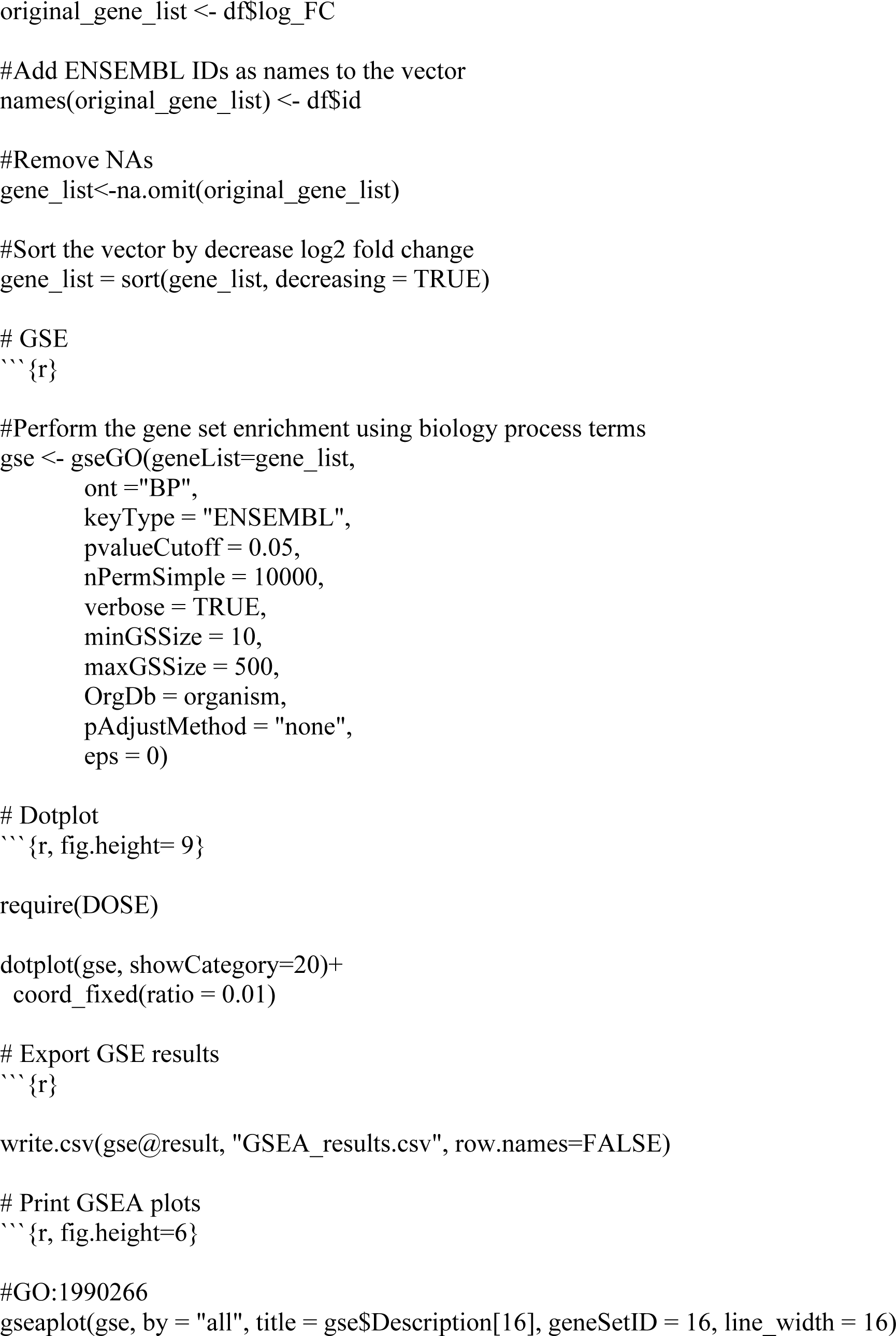
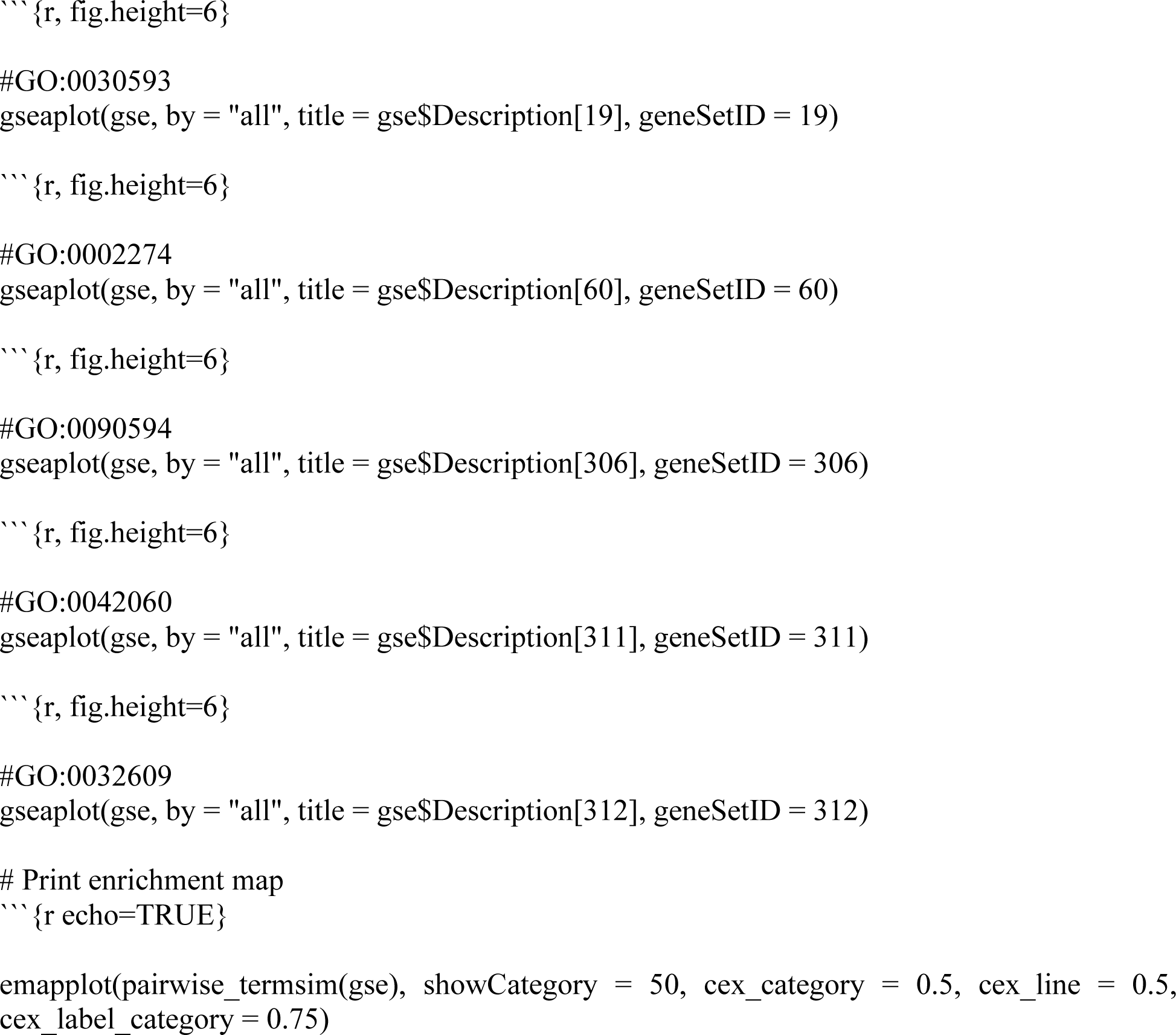

