## Supplemental figures for "A type 1 immune-stromal cell network mediates disease tolerance and barrier protection against intestinal infection"

**Figure S1.**

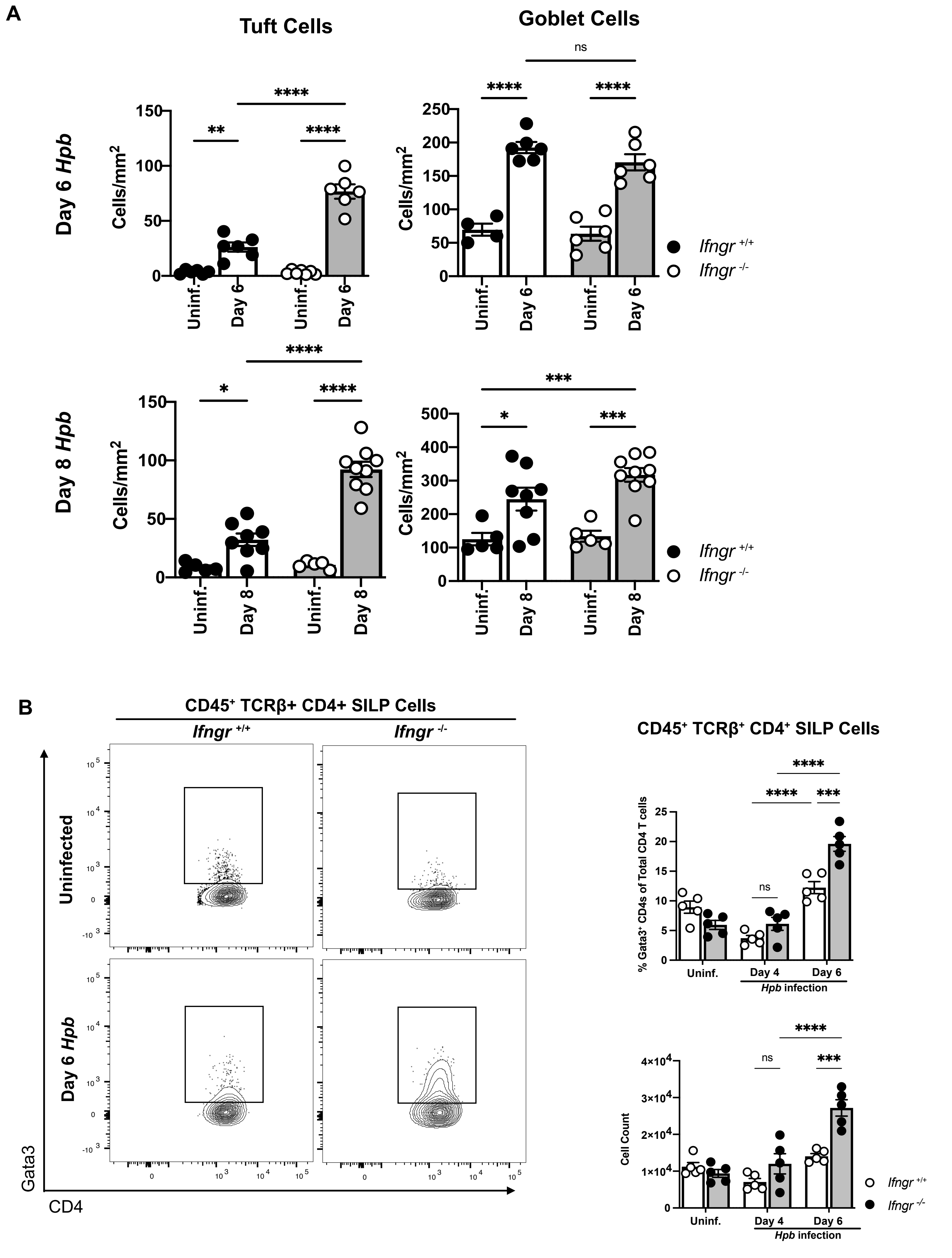
A
Tuft Cells
Goblet Cells
Day 6 Hpb
Day 8 Hpb
Cells/mm2
Uninf.
Day 6
Day 8
ns
Ifngr +/+
Ifngr -/-
B
CD45+ TCRβ+ CD4+ SILP Cells
Ifngr +/+
Ifngr -/-
Uninfected
Day 6 Hpb
Gata3
CD4
CD45+ TCRβ+ CD4+ SILP Cells
% Gata3+ CD4s of Total CD4 T cells
Cell Count
Uninf.
Day 4
Day 6
Hpb infection
ns
Ifngr +/+
Ifngr -/-

**Figure S2.**

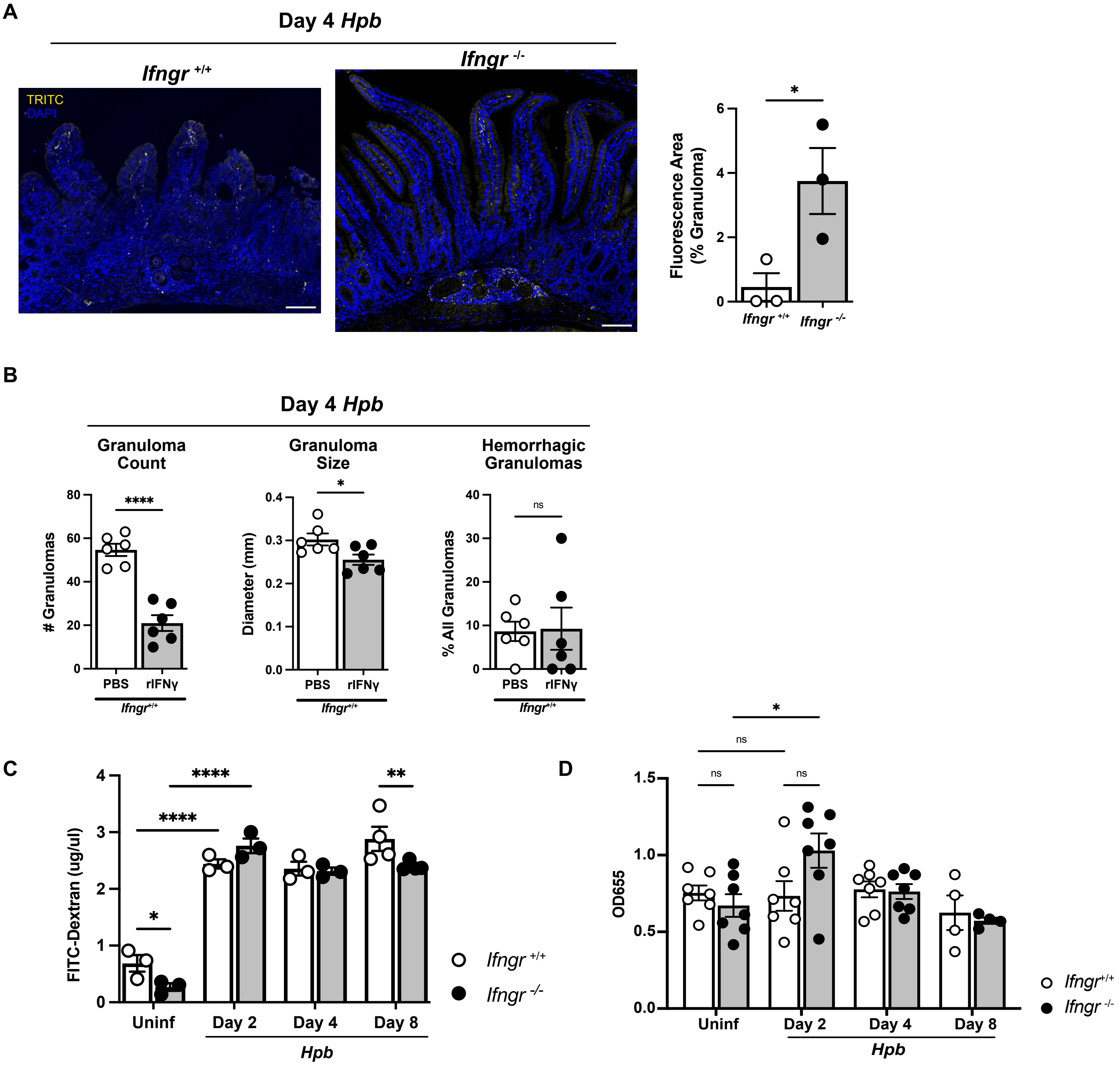
A
Day 4 Hpb
Ifngr +/+
Ifngr -/-
TRITC
DAPI
Fluorescence Area (% Granuloma)
*
0
2
4
6
Ifngr +/+
Ifngr -/-
B
Day 4 Hpb
Granuloma Count
Granuloma Size
Hemorrhagic Granulomas
### Granulomas
****
0
20
40
60
80
PBS
rIFNγ
Ifngr+/+
Diameter (mm)
*
0.0
0.1
0.2
0.3
0.4
PBS
rIFNγ
Ifngr+/+
% All Granulomas
ns
0
10
20
30
40
PBS
rIFNγ
Ifngr+/+
C
FITC-Dextran (ug/ul)
0
1
2
3
4
*
****
****
**
Uninf
Day 2
Day 4
Day 8
Hpb
Ifngr +/+
Ifngr -/-
D
OD655
0.0
0.5
1.0
1.5
*
ns
ns
ns
Uninf
Day 2
Day 4
Day 8
Hpb
Ifngr+/+
Ifngr -/-

**Figure S3.**

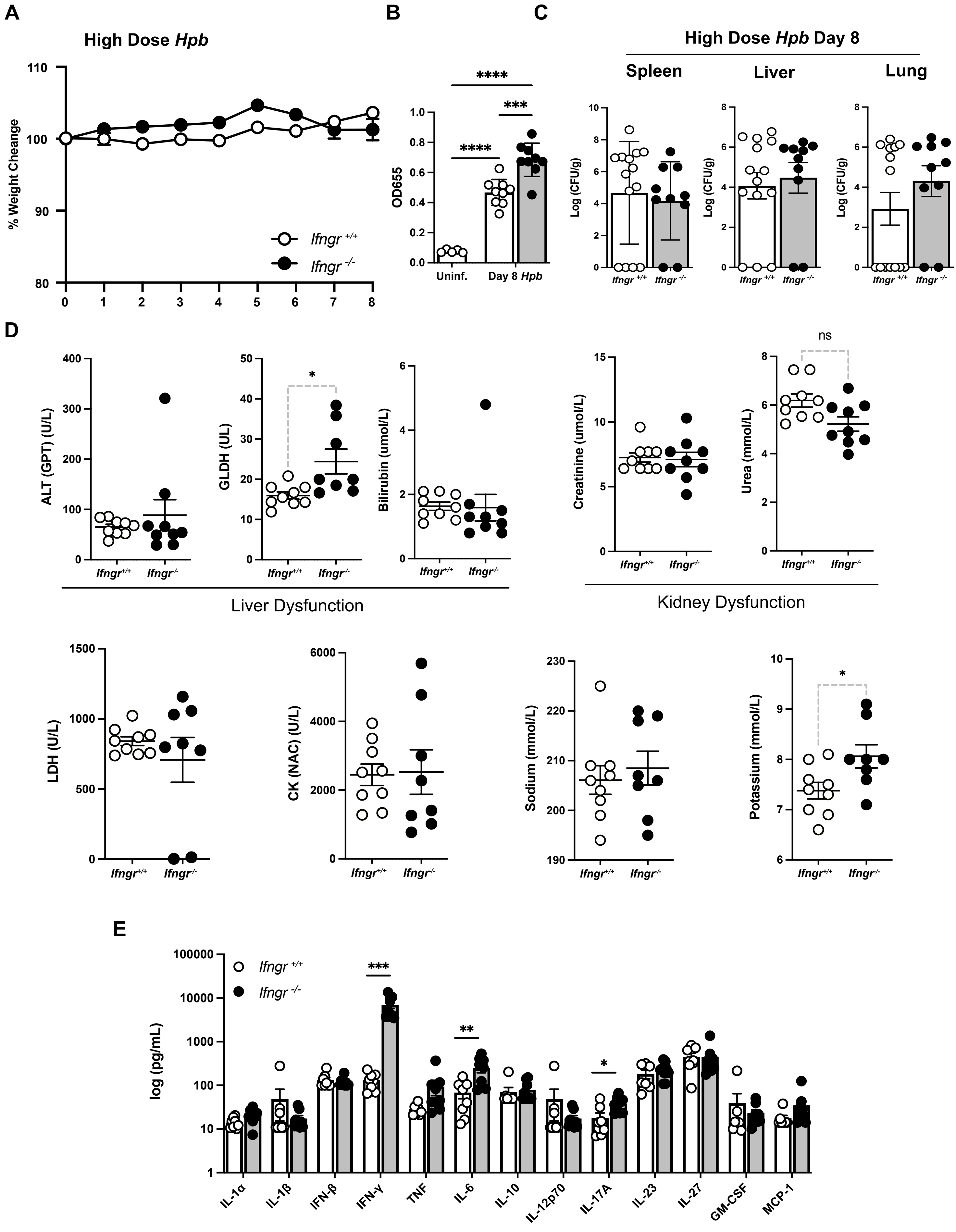
A
B
C
High Dose Hpb
% Weight Cheange
Ifngr +/+
Ifngr -/-
OD655
Uninf.
Day 8 Hpb
****
***
High Dose Hpb Day 8
Spleen
Liver
Lung
Log (CFU/g)
D
ALT (GPT) (U/L)
GLDH (UL)
Bilirubin (umol/L)
Creatinine (umol/L)
Urea (mmol/L)
ns
Liver Dysfunction
Kidney Dysfunction
LDH (U/L)
CK (NAC) (U/L)
Sodium (mmol/L)
Potassium (mmol/L)
E
log (pg/mL)
IL-1α
IL-1β
IFN-β
IFN-γ
TNF
IL-6
IL-10
IL-12p70
IL-17A
IL-23
IL-27
GM-CSF
MCP-1

**Figure S4.**

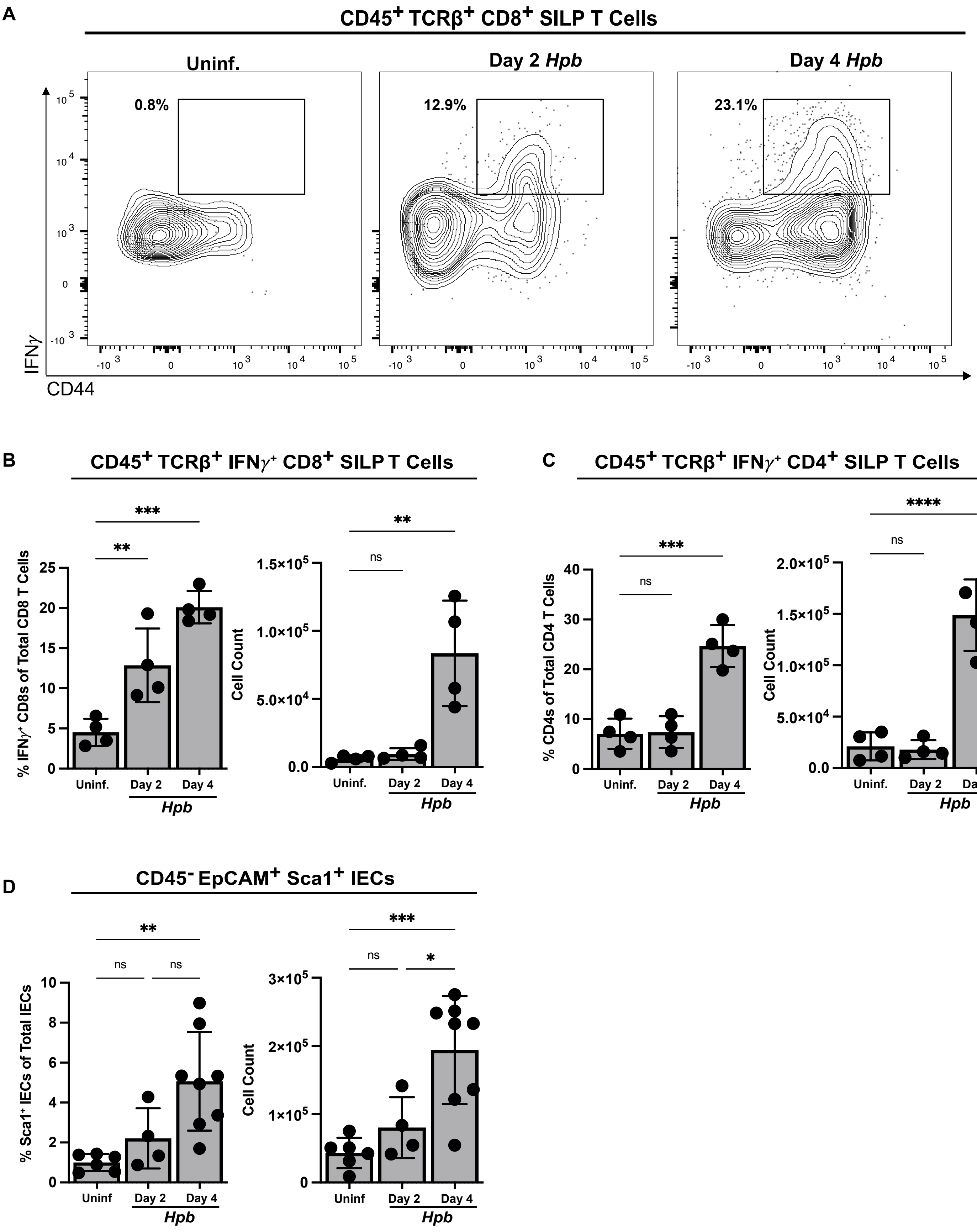
A
CD45+ TCRβ+ CD8+ SILP T Cells
Uninf.
Day 2 Hpb
Day 4 Hpb
0.8%
12.9%
23.1%
IFNγ
CD44
B
CD45+ TCRβ+ IFNγ+ CD8+ SILP T Cells
% IFNγ+ CD8s of Total CD8 T Cells
Cell Count
Uninf.
Day 2
Day 4
Hpb
C
CD45+ TCRβ+ IFNγ+ CD4+ SILP T Cells
% CD4s of Total CD4 T Cells
Cell Count
D
CD45- EpCAM+ Sca1+ IECs
% Sca1+ IECs of Total IECs
Cell Count
Uninf
ns

**Figure S5.**

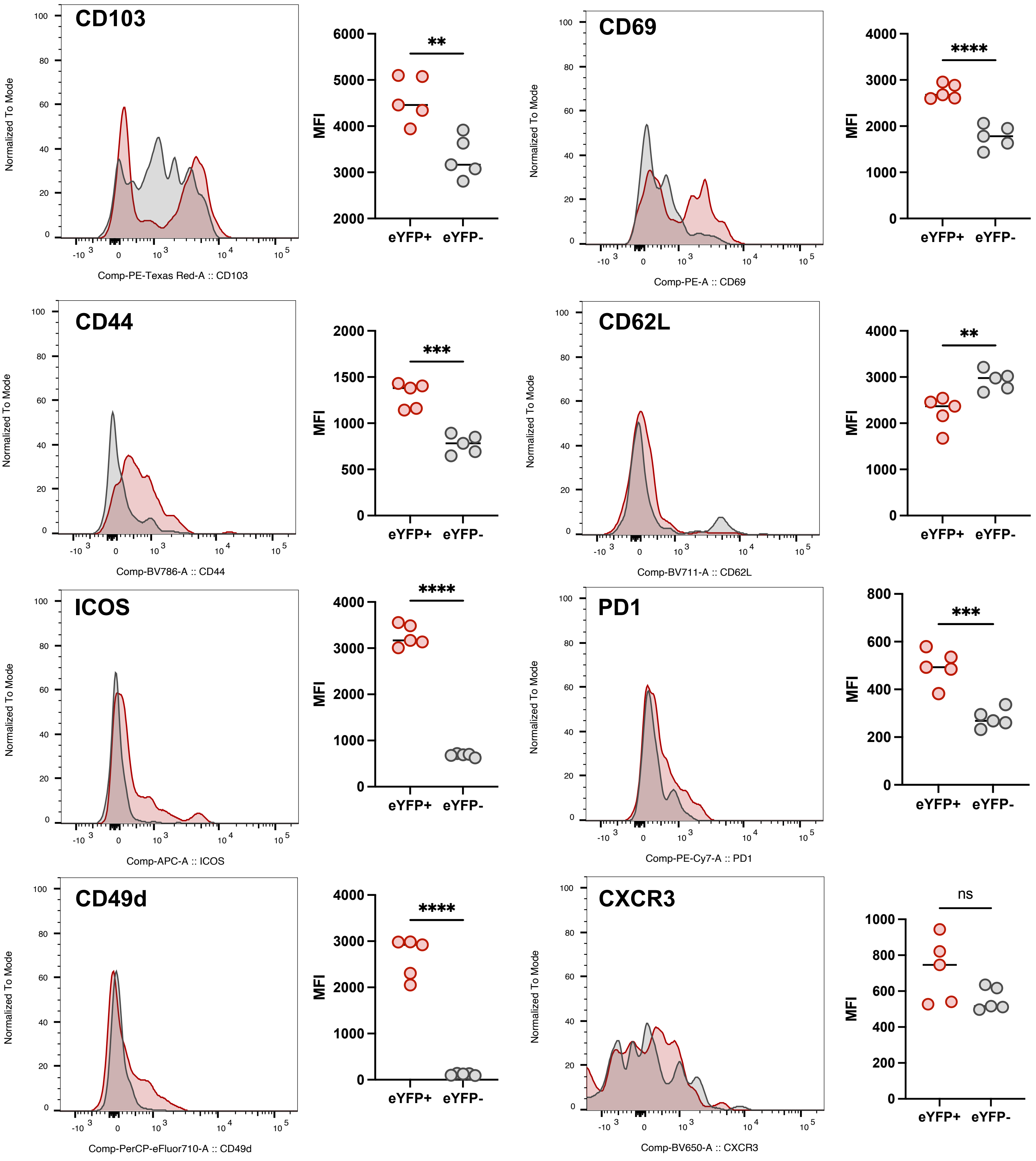
CD103
Normalized To Mode
Comp-PE-Texas Red-A :: CD103
MFI
**
eYFP+
eYFP-
CD69
Comp-PE-A :: CD69
****
CD44
Comp-BV786-A :: CD44
***
CD62L
Comp-BV711-A :: CD62L
**
ICOS
Comp-APC-A :: ICOS
****
PD1
Comp-PE-Cy7-A :: PD1
***
CD49d
Comp-PerCP-eFluor710-A :: CD49d
****
CXCR3
Comp-BV650-A :: CXCR3
ns

**Figure S6.**

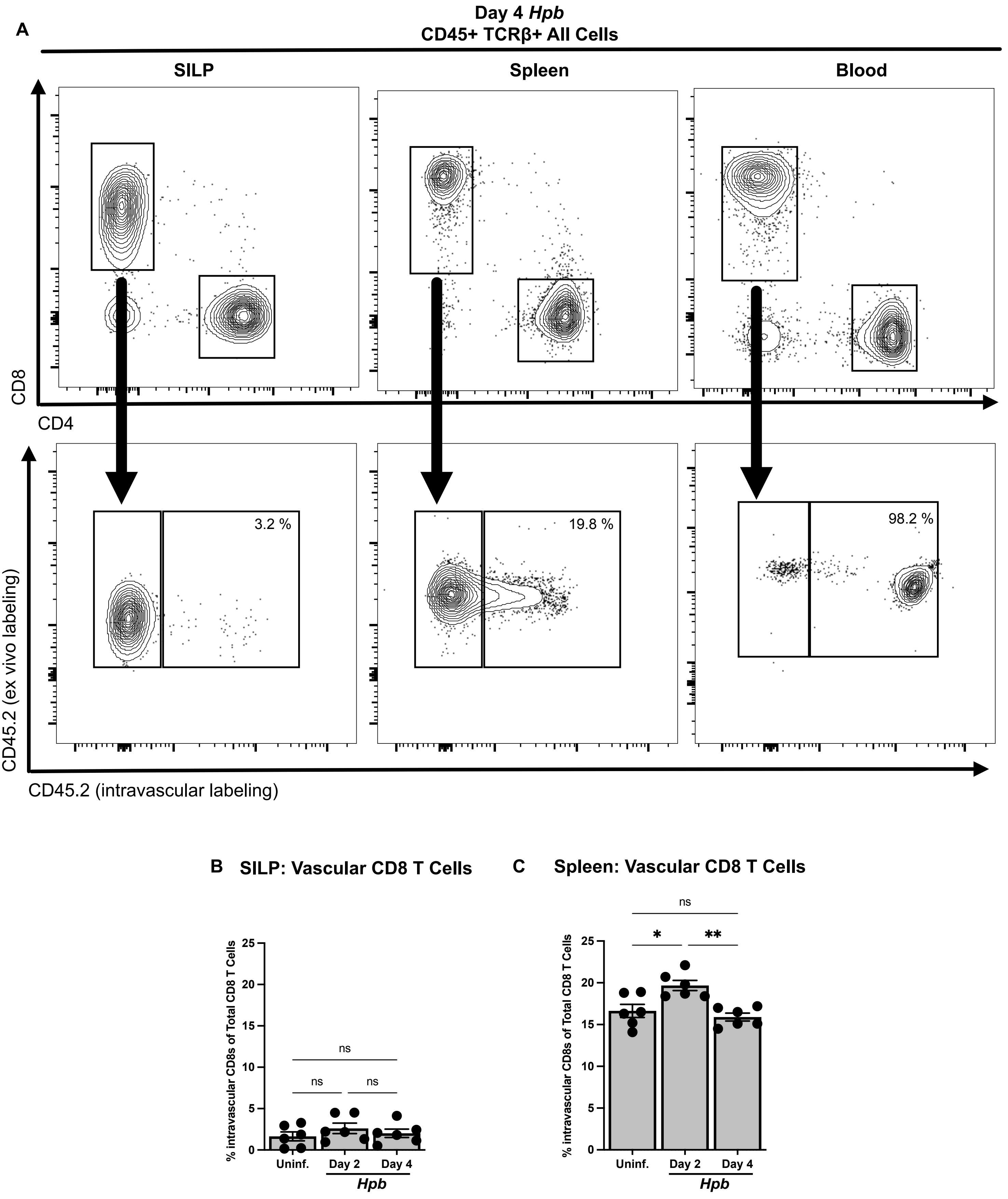
A
Day 4 Hpb
CD45+ TCRβ+ All Cells
SILP
Spleen
Blood
CD8
CD4
3.2 %
19.8 %
98.2 %
CD45.2 (ex vivo labeling)
CD45.2 (intravascular labeling)
B
SILP: Vascular CD8 T Cells
% intravascular CD8s of Total CD8 T Cells
ns
ns
ns
Uninf.
Day 2
Day 4
Hpb
C
Spleen: Vascular CD8 T Cells
% intravascular CD8s of Total CD8 T Cells
ns
*
**
Uninf.
Day 2
Day 4
Hpb

**Figure S7.**

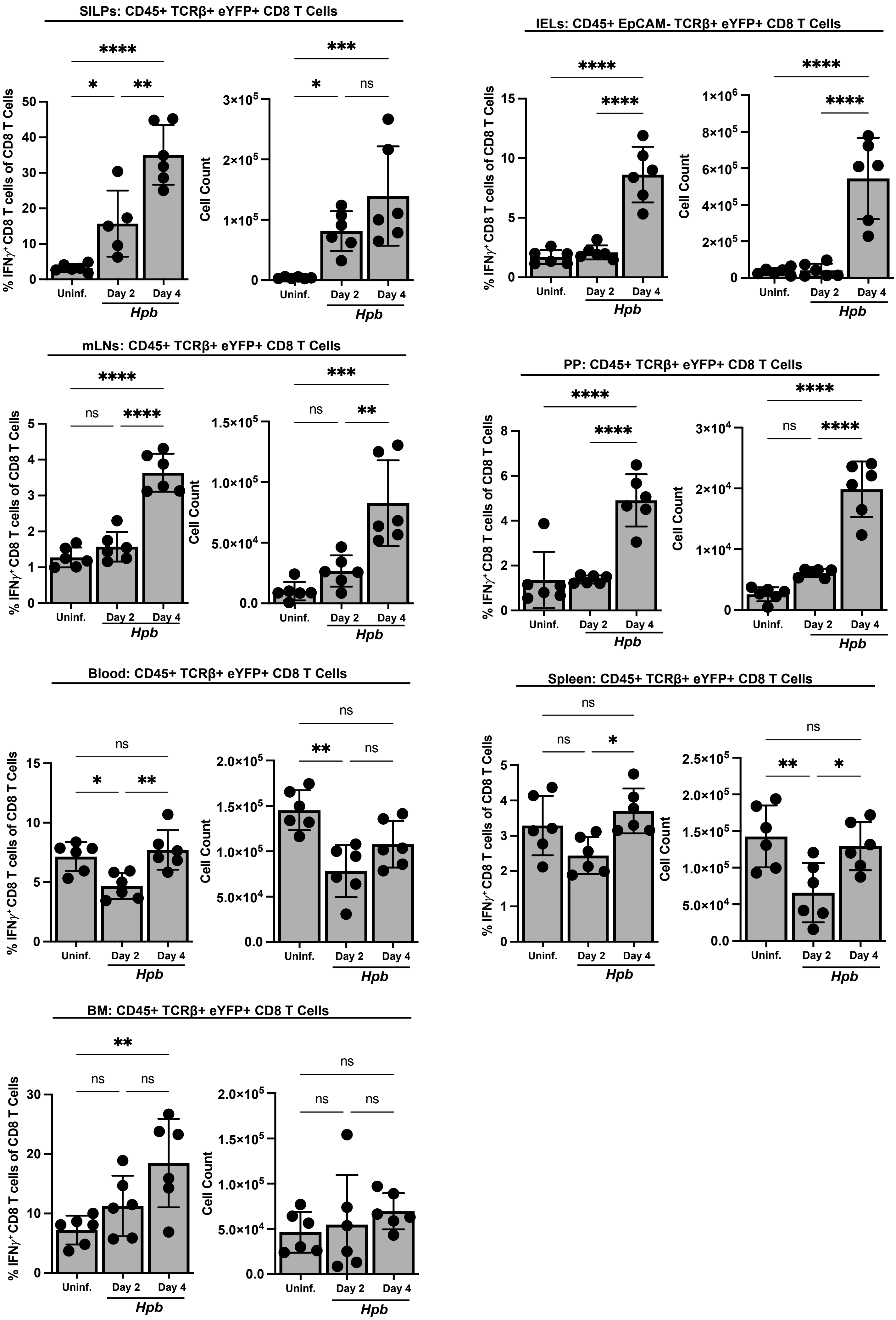
SILPs: CD45+ TCRβ+ eYFP+ CD8 T Cells
IELs: CD45+ EpCAM- TCRβ+ eYFP+ CD8 T Cells
mLNs: CD45+ TCRβ+ eYFP+ CD8 T Cells
PP: CD45+ TCRβ+ eYFP+ CD8 T Cells
Blood: CD45+ TCRβ+ eYFP+ CD8 T Cells
Spleen: CD45+ TCRβ+ eYFP+ CD8 T Cells
BM: CD45+ TCRβ+ eYFP+ CD8 T Cells
% IFNγ+ CD8 T cells of CD8 T Cells
Cell Count
Uninf.
Day 2
Day 4
Hpb
ns
*
**
***
****

**Figure S8.**

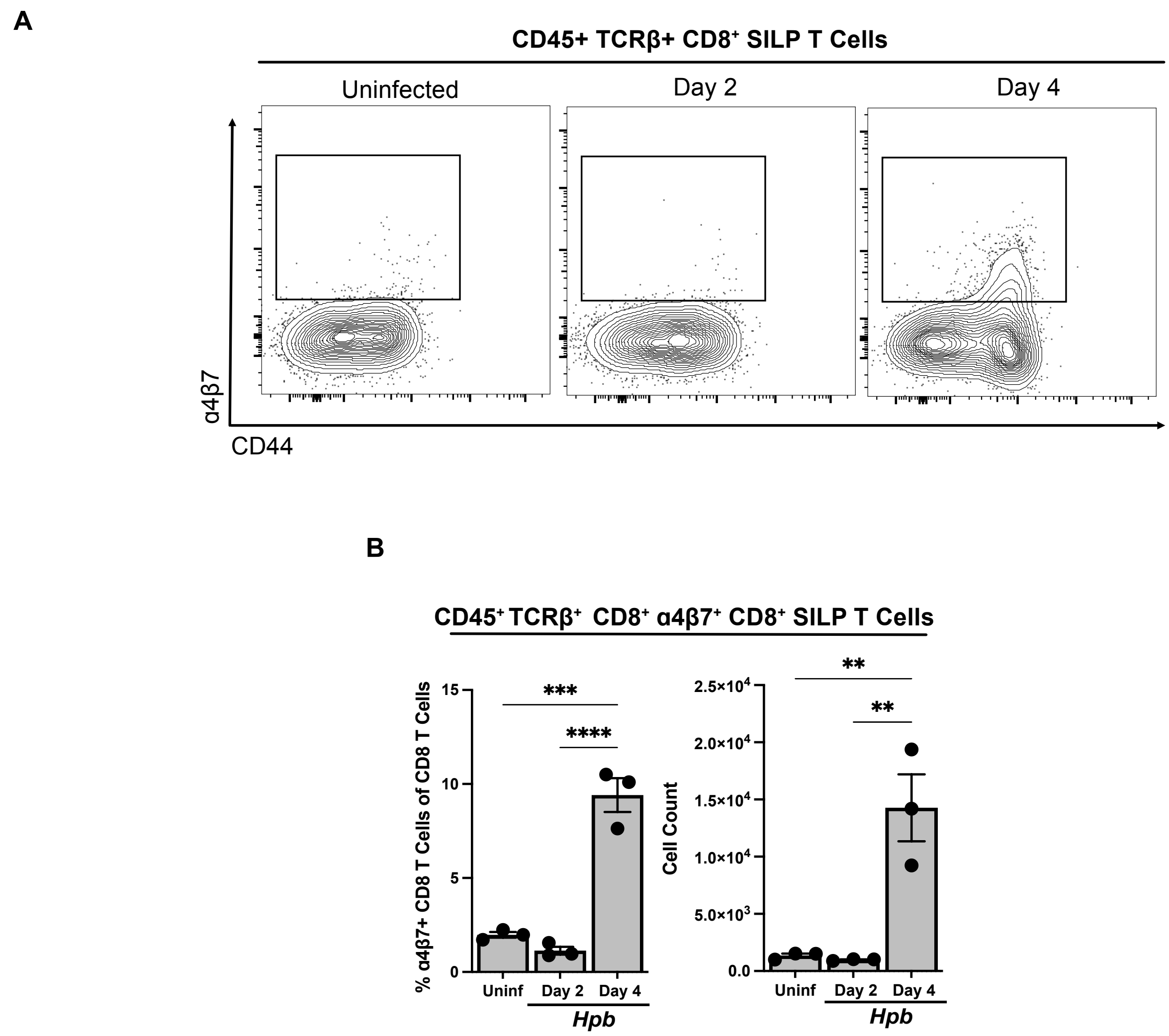
A
CD45+ TCRβ+ CD8+ SILP T Cells
Uninfected
Day 2
Day 4
α4β7
CD44
B
CD45+ TCRβ+ CD8+ α4β7+ CD8+ SILP T Cells
% α4β7+ CD8 T Cells of CD8 T Cells
15
10
5
0
***
****
Uninf
Day 2
Day 4
Hpb
Cell Count
2.5×10^4
2.0×10^4
1.5×10^4
1.0×10^4
5.0×10^3
0.0
**
**
Uninf
Day 2
Day 4
Hpb

#### Figure S9.

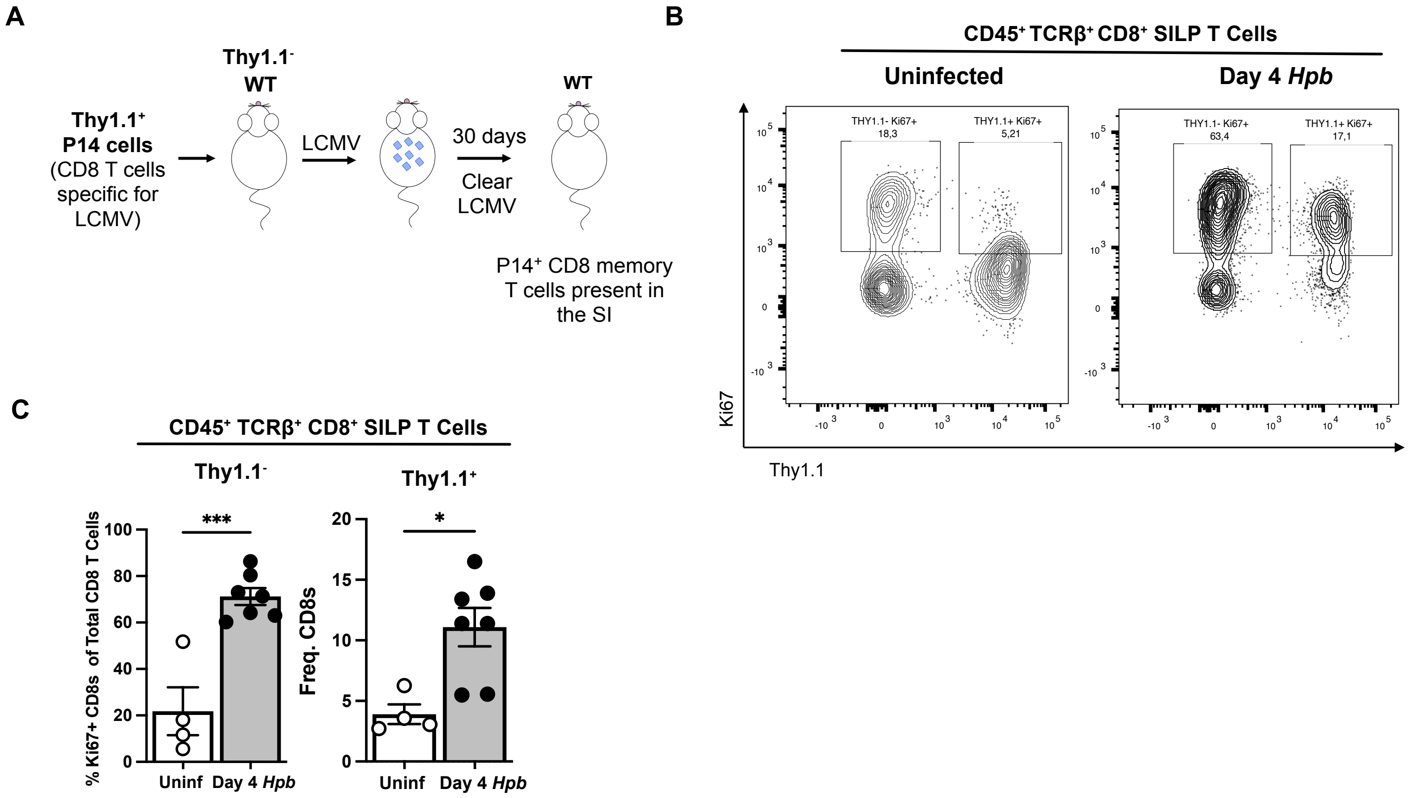
A
Thy1.1+
P14 cells
(CD8 T cells specific for LCMV)
Thy1.1-
WT
LCMV
30 days
Clear LCMV
WT
P14+ CD8 memory T cells present in the SI
B
CD45+ TCRβ+ CD8+ SILP T Cells
Uninfected
Day 4 Hpb
THY1.1- Ki67+
18,3
THY1.1+ Ki67+
5,21
THY1.1- Ki67+
63,4
THY1.1+ Ki67+
17,1
Ki67
Thy1.1
C
CD45+ TCRβ+ CD8+ SILP T Cells
Thy1.1-
Thy1.1+
% Ki67+ CD8s of Total CD8 T Cells
Freq. CD8s
***
*
Uninf
Day 4 Hpb

**Figure S10.**

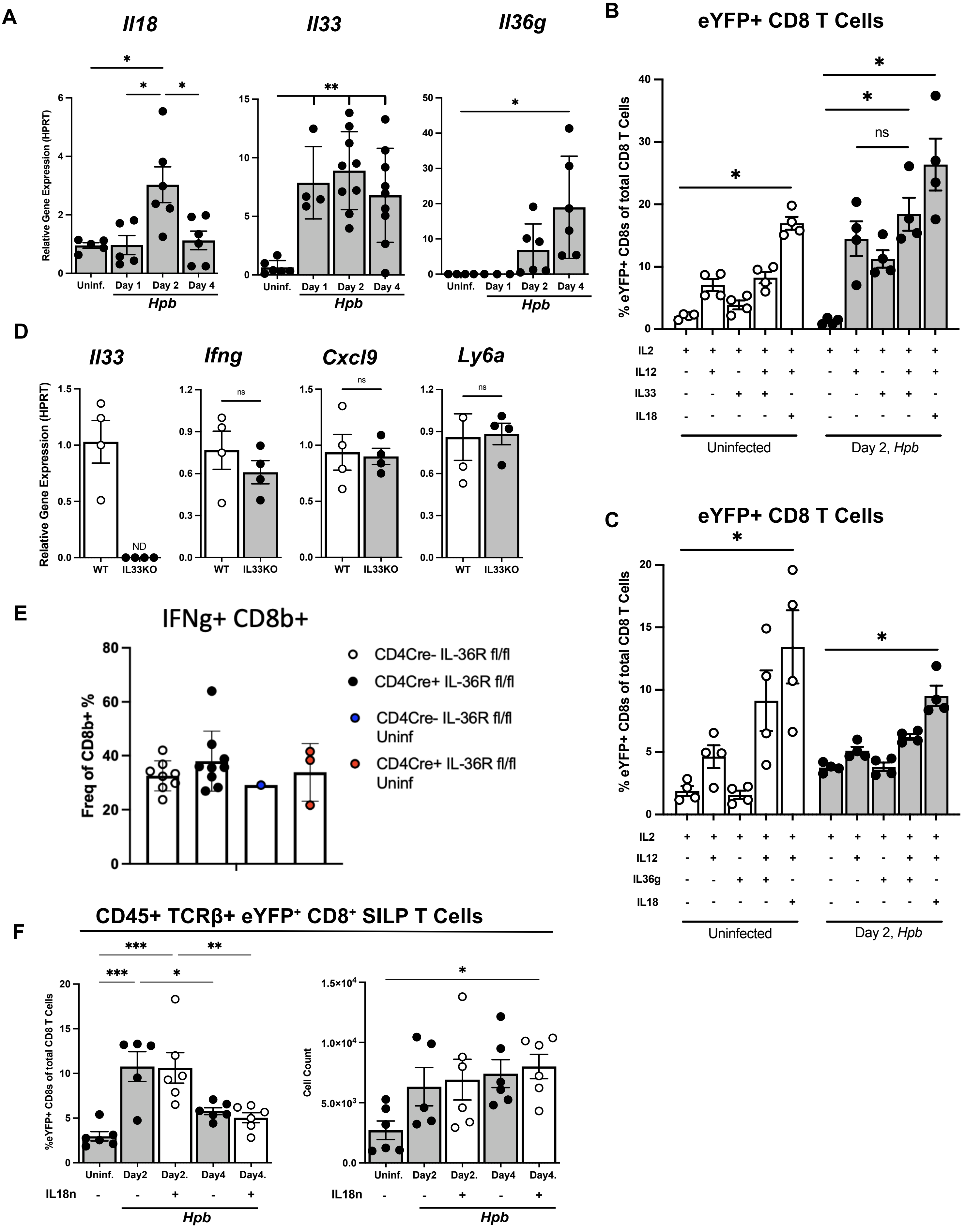
A
Il18
Il33
Il36g
Relative Gene Expression (HPRT)
Uninf.
Day 1
Day 2
Day 4
Hpb
B
eYFP+ CD8 T Cells
% eYFP+ CD8s of total CD8 T Cells
IL2
IL12
IL33
IL18
Uninfected
Day 2, Hpb
ns
C
eYFP+ CD8 T Cells
% eYFP+ CD8s of total CD8 T Cells
IL2
IL12
IL36g
IL18
Uninfected
Day 2, Hpb
D
Il33
Ifng
Cxcl9
Ly6a
Relative Gene Expression (HPRT)
ND
ns
WT
IL33KO
E
IFNg+ CD8b+
Freq of CD8b+ %
CD4Cre- IL-36R fl/fl
CD4Cre+ IL-36R fl/fl
CD4Cre- IL-36R fl/fl Uninf
CD4Cre+ IL-36R fl/fl Uninf
F
CD45+ TCRβ+ eYFP+ CD8+ SILP T Cells
%eYFP+ CD8s of total CD8 T Cells
Cell Count
Uninf.
Day2
Day2.
Day4
Day4.
IL18n
Hpb

### Figure S11.

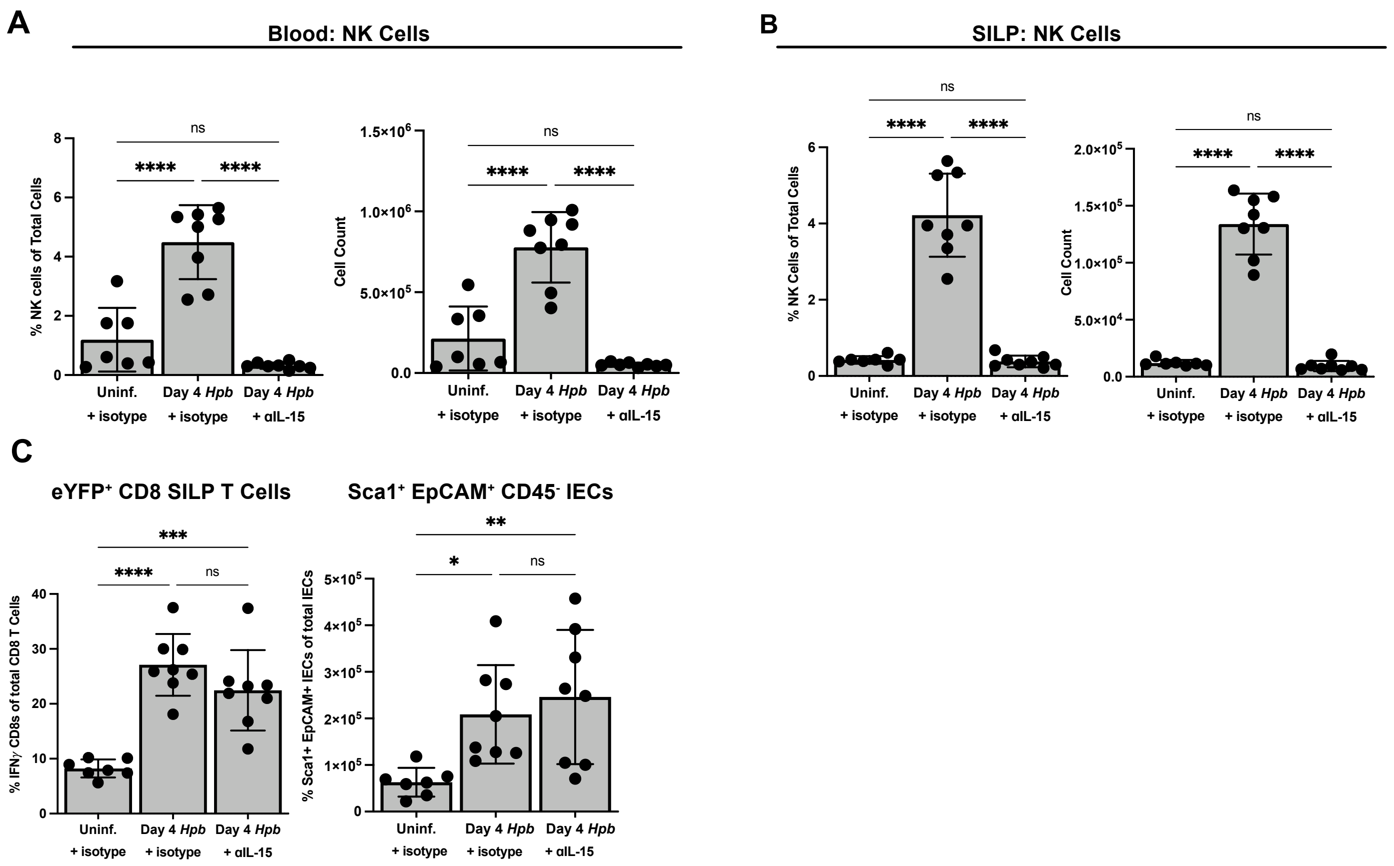
A
Blood: NK Cells
% NK cells of Total Cells
Cell Count
ns
****
1.5×10^6
1.0×10^6
5.0×10^5
0.0
Uninf.
+ isotype
Day 4 Hpb
+ isotype
Day 4 Hpb
+ αIL-15
B
SILP: NK Cells
% NK Cells of Total Cells
2.0×10^5
1.5×10^5
1.0×10^5
5.0×10^4
C
eYFP+ CD8 SILP T Cells
Sca1+ EpCAM+ CD45- IECs
% IFNγ CD8s of total CD8 T Cells
% Sca1+ EpCAM+ IECs of total IECs
***
**
*
5×10^5
4×10^5
3×10^5
2×10^5
1×10^5

**Figure S12.**

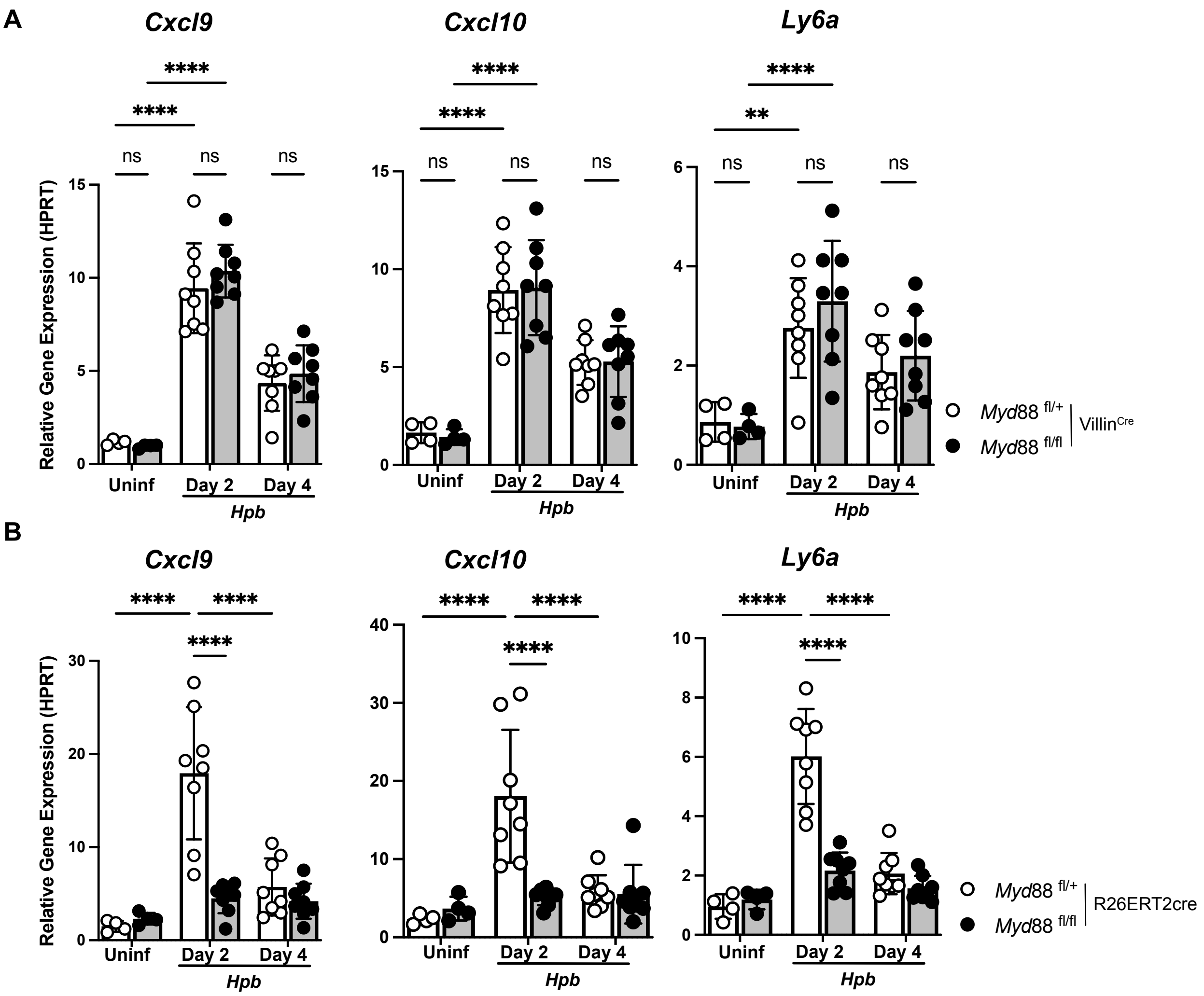
A
Cxcl9
Cxcl10
Ly6a
****
**
ns
Relative Gene Expression (HPRT)
Uninf
Day 2
Day 4
Hpb
Myd88 fl/+
Myd88 fl/fl
VillinCre
B
Cxcl9
Cxcl10
Ly6a
Myd88 fl/+
Myd88 fl/fl
R26ERT2cre

**Figure S13.**

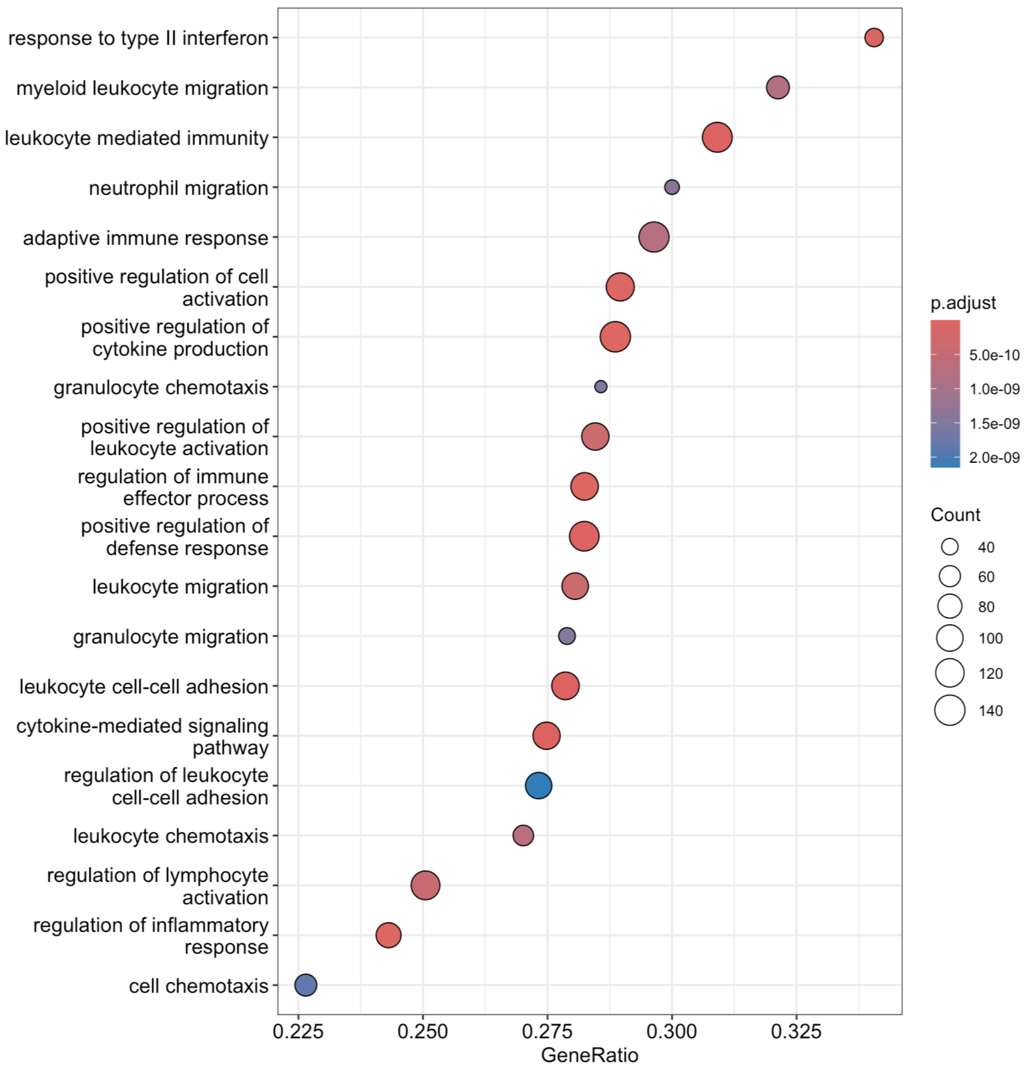

response to type II interferon
myeloid leukocyte migration
leukocyte mediated immunity
neutrophil migration
adaptive immune response
positive regulation of cell activation
positive regulation of cytokine production
granulocyte chemotaxis
positive regulation of leukocyte activation
regulation of immune effector process
positive regulation of defense response
leukocyte migration
granulocyte migration
leukocyte cell-cell adhesion
cytokine-mediated signaling pathway
regulation of leukocyte cell-cell adhesion
leukocyte chemotaxis
regulation of lymphocyte activation
regulation of inflammatory response
cell chemotaxis
0.225
0.250
0.275
0.300
0.325
GeneRatio
p.adjust
5.0e-10
1.0e-09
1.5e-09
2.0e-09
Count
40
60
80
100
120
140

**Figure S14.**

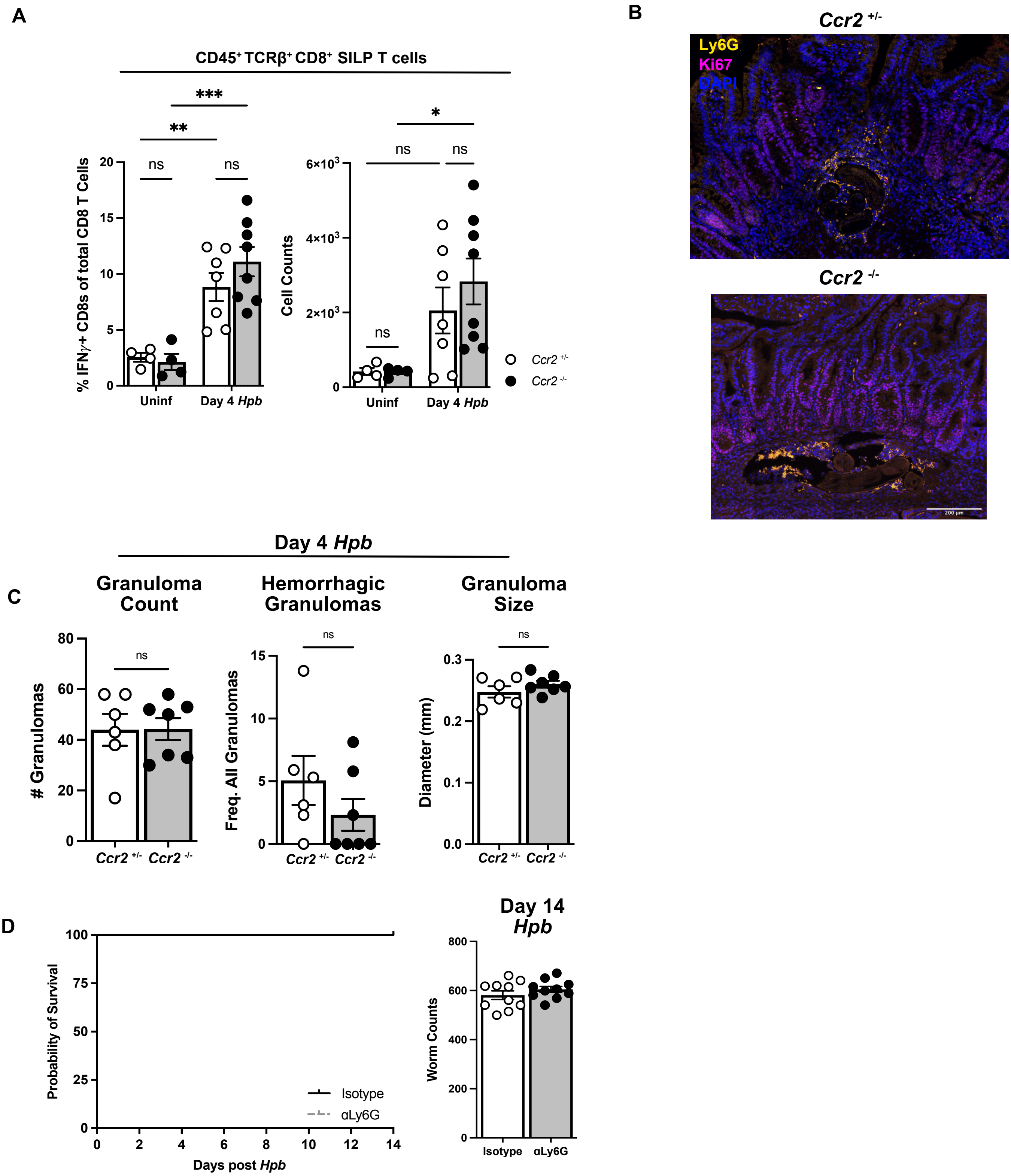

A
CD45+ TCRβ+ CD8+ SILP T cells
% IFNγ+ CD8s of total CD8 T Cells
Cell Counts
Uninf
Day 4 Hpb
ns
**
***
*
Ccr2 +/-
Ccr2 -/-
B
Ccr2 +/-
Ccr2 -/-
Ly6G
Ki67
DAPI
Day 4 Hpb
C
Granuloma Count
Hemorrhagic Granulomas
Granuloma Size
### Granulomas
Freq. All Granulomas
Diameter (mm)
D
Probability of Survival
Days post Hpb
Isotype
αLy6G
Day 14 Hpb
Worm Counts

Figure S15.

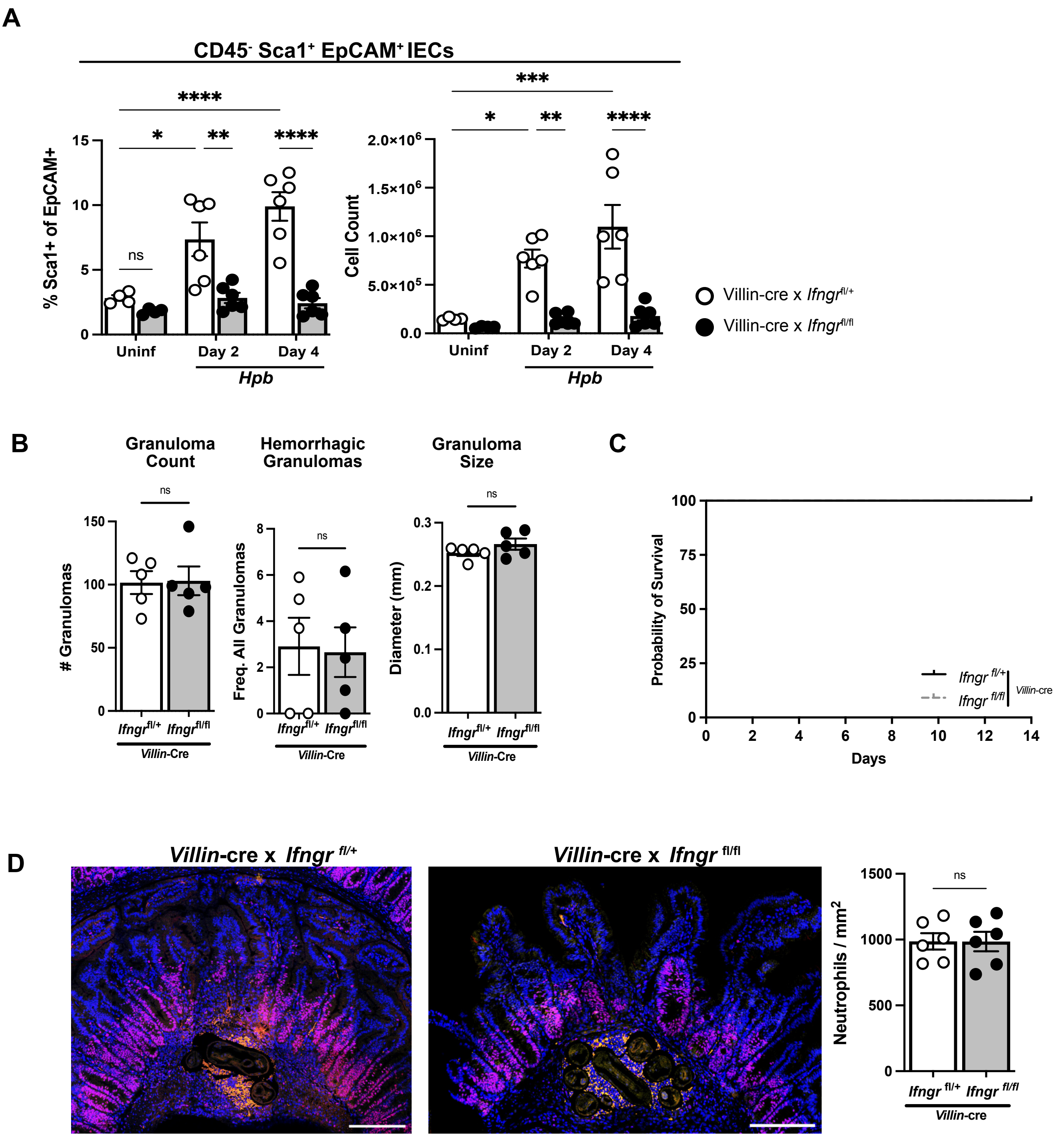

A
CD45- Sca1+ EpCAM+ IECs
% Sca1+ of EpCAM+
Cell Count
Uninf
Day 2
Day 4
Hpb
ns
Villin-cre x Ifngr fl/+
Villin-cre x Ifngr fl/fl
B
Granuloma Count
Hemorrhagic Granulomas
Granuloma Size
### Granulomas
Freq. All Granulomas
Diameter (mm)
Ifngr fl/+
Ifngr fl/fl
Villin-Cre
C
Probability of Survival
Days
Villin-cre
D
Villin-cre x Ifngr fl/+
Villin-cre x Ifngr fl/fl
Neutrophils / mm2

**Figure S16.**

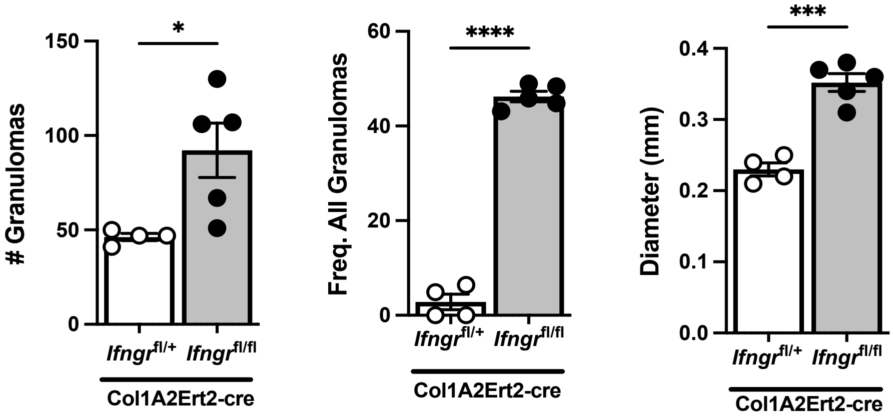

150
100
50
0
### Granulomas
*
Ifngr fl/+
Ifngr fl/fl
Col1A2Ert2-cre
60
40
20
0
Freq. All Granulomas
****
Ifngr fl/+
Ifngr fl/fl
Col1A2Ert2-cre
0.4
0.3
0.2
0.1
0.0
Diameter (mm)
***
Ifngr fl/+
Ifngr fl/fl
Col1A2Ert2-cre
